# Transposon targeting non-coding RNA transcription targets G/C-rich tracts and is facilitated by an intrinsically disordered protein in *Tetrahymena*

**DOI:** 10.1101/2025.10.09.681302

**Authors:** Xia Cai, Yike Liu, Jialuo Li, Zhuoran Zhao, Xinyu Yao, Zhihao Zhai, Mingmei Liucong, Yujie Liu, Xue Wang, Kazufumi Mochizuki, Miao Tian

**Author notes:** Corresponding author: Miao Tian. Contributed equally to this work (Xia Cai, Yike Liu).

## Abstract

*Tetrahymena* is a well-established ciliate model organism, known for its nuclear dualism and unique mechanism for transposon defense, wherein transposons are kept within the highly heterochromatinized micronucleus (MIC), which is transcriptionally inert except during meiosis. The discovery of preferential transcription in the meiotic MIC at regions enriched with transposons, and the identification of MIC transcription regulatory proteins, emphasizes the necessity to elucidate the underlying mechanisms driving this process. In this study, we demonstrate that G/C-rich tracts are overrepresented in meiotic non-coding RNA (ncRNA) transcription regions and that RNA polymerase II (Pol II) is highly enriched at and around these tracts. Further analysis revealed that Pol II’s association with G/C-rich tracts is not abolished in cells lacking Rib1, a Mediator complex-associated protein essential for MIC ncRNA biogenesis. Nevertheless, in the absence of Rib1, Pol II showed abnormal association with the meiotic MIC chromatin, suggesting that Rib1 is critical for maintaining Pol II’s binding specificity. Through Rib1 truncation analysis, we found that the intrinsically disordered region, which contains putative phase-separating peptides, is crucial for its function. Disruption of phase-separation, either by deleting these peptides or by treating cells with a phase-separation disruption reagent, leads to aberrant localization of both Rib1 and Pol II on the MIC chromatin, implicating that Rib1 likely facilitates the MIC ncRNA transcription via phase-separation.

## Introduction

Ciliates are single-celled eukaryotes that typically host two functionally distinct types of nuclei. The smaller nucleus is called the micronucleus (MIC), and it contains the germline genome, while the larger nucleus is termed the macronucleus (MAC). The MAC is derived from the MIC, and it functions as a somatic nucleus. Although the MIC genome contains abundant transposable elements (TEs), TE remnants, and other repetitive sequences like most of other eukaryotic genomes, the MAC genome is nearly devoid of these sequences (see (Hamilton et al., 2016; Jin et al., 2023; Wells and Feschotte, 2020)). This is because TEs are eliminated as internal eliminate sequences (IESs) from the MAC destined zygotic nucleus (hereinafter referred to as the MAC anlagen) (see (Jiang et al., 2025; Long et al., 2023; Orias et al., 2011; Ruehle et al., 2016)). Ciliates use the streamlined TE-free MAC genome for transcription of mRNAs as well as functional non-coding RNAs (ncRNAs), such as ribosomal RNAs (rRNAs, (Leer et al., 1979)). In contrast, the MIC chromatin is more compact than their MAC chromatin and transcriptionally inactive in most life stages (Raikhel et al., 1981; Raikov et al., 1989; Skarlato, 1982; Wolfe et al., 1976). The inheritance of the germline genome in the transcriptionally-inactive MIC with compact chromatin conformation is believed to be beneficial for keeping the genome integrity (see (Drews et al., 2022; Maeshima et al., 2021)). Therefore, the binucleated ciliates showcase a unique strategy for repressing TEs: by eliminating them from the transcriptionally active nucleus, while keeping them in another transcriptionally inert nucleus. Given that the MAC analgen originates from the MIC, which contains IESs, a compelling question arises: how do cells specifically eliminate IESs from the zygotic genome to generate a TE-free MAC genome?

The exceptional transcription in the MIC at the onset of sexual reproduction stage (Cai et al., 2025; Gruchota et al., 2017; Mochizuki and Gorovsky, 2004; Owsian et al., 2022; Tian et al., 2022) plays an important role in regulating the specific IES elimination in some model ciliates, such as *Tetrahymena thermophila* and *Paramecium tetraurelia* (see (Allen and Nowacki, 2020; Chalker et al., 2013; Drews et al., 2022; Gao et al., 2023; Lyu et al., 2024; Noto and Mochizuki, 2017)). Taking *Tetrahymena* as an example, ncRNAs are produced by the otherwise silent MIC during meiotic prophase. Because the MIC lacks the factors necessary to process nascent RNAs into mature messenger RNAs (mRNAs), its transcripts are not translated into proteins (Zhao et al., 2019). The MIC ncRNA transcription takes place preferentially at and around a subset of TEs and their remnants, also known as the Type A IESs (Noto et al., 2015). Likely due to the bidirectional transcriptional nature of the MIC genome (Chalker and Yao, 2001; Schoeberl et al., 2012), these ncRNAs form double-stranded RNAs (perhaps via annealing), which then serve as substrates for a nuclear Dicer-like protein, Dcl1 (Malone et al., 2005; Mochizuki and Gorovsky, 2005). The cleavage by Dcl1 produces ∼27–31 nucleotide small RNAs, referred to as scan RNAs (scnRNAs) (Lee and Collins, 2006; Mochizuki and Kurth, 2013), which are incorporated with a *Tetrahymena* Piwi protein, Twi1, forming Twi1-scnRNA complexes (Mochizuki et al., 2002). Then, these complexes are transported to the parental MAC (pMAC) (Noto et al., 2010), where they interact with the pMAC long non-coding RNAs (lncRNAs). This interaction leads to the clearance of scnRNAs that are complementary to the MAC sequences (i.e., non-TEs, (Aronica et al., 2008; Noto and Mochizuki, 2018; Shehzada et al., 2024)), while resulting in a pool of Twi1-scnRNAs that are highly specific to IESs (Noto et al., 2015) (Figure 1). These retained scnRNAs are subsequently transported into the new MAC anlagen, where they trigger the initiation of the second round of scnRNA biogenesis from IESs (Noto et al., 2015). Ultimately, these scnRNAs direct the targeted elimination of IESs (Cheng et al., 2010; Liu et al., 2007; Taverna et al., 2002; Vogt and Mochizuki, 2013; Zhao et al., 2019).

**Figure 1.**
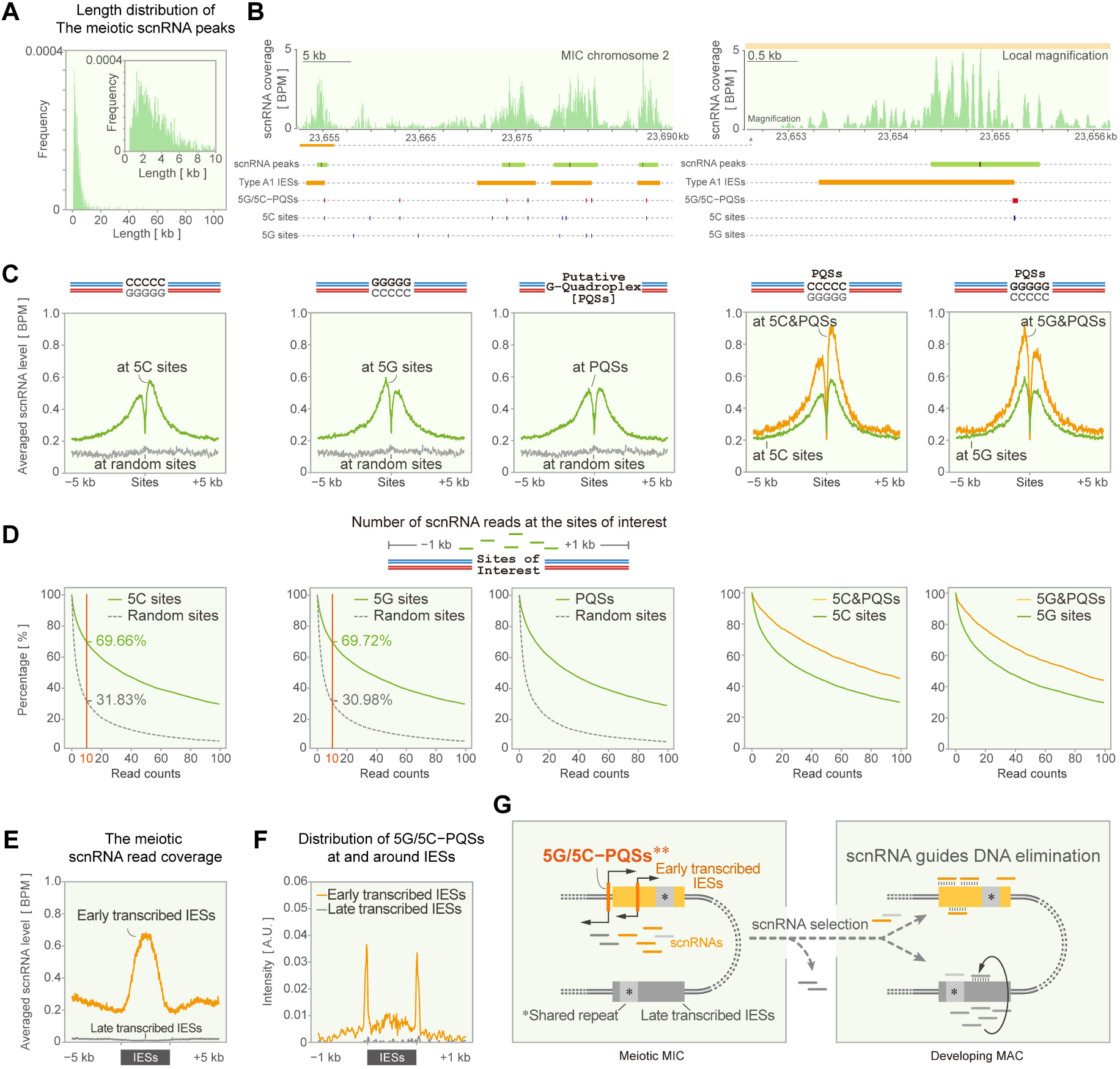
Putative quadruplex-forming sequences containing 5G or 5C tracts are positively correlated with the meiotic MIC transcription. A, Distribution of the meiotic scnRNA peak lengths. B, Left panel: A representative genomic region (CM028433.1:23,652,282-23,690,678) illustrating the reliable identification of scnRNA peaks (green horizontal bars) and their corresponding peak summits (black vertical lines). Additionally, Type A1 IESs (orange bars), 5G or 5C tract-containing putative DNA-quadruplex formation sequences (5G/5C-PQSs, red vertical lines), and 5G or 5C tracts (blue vertical lines) are also displayed. The scnRNA sequence read coverage is normalized using bins per million mapped reads (BPM). Right panel: A magnified view of a specific genomic locus depicted in the left panel (marked with an orange bar). C, Averaged WT meiotic scnRNA sequencing read coverage at and surrounding sites of interest. D, Occurrence frequency of WT meiotic scnRNAs from regions containing specific features. For comparison, equal numbers of random regions were also analyzed. E, Averaged WT meiotic scnRNA sequencing read distribution at and surrounding Type A1 and Type B2 IESs. F, Distribution of 5G/5C-PQSs at and surrounding Type A1 and Type B2 IESs. G, Schematic illustration of meiotic ncRNA transcription, which preferentially occurs at early transcribed IESs containing 5G/5C-PQSs. After selection, scnRNAs derived from non-IES regions are removed, while those derived from early-transcribed IESs are transported to the developing MAC, where they stimulate the transcription of late-transcribed IESs. Ultimately, these scnRNAs guide IES elimination. “*” denotes repetitive sequences shared by early and late transcribed IESs. “**” not all IESs have these motifs.

The characterization of the DNA elimination mechanism once revealed unanticipated essential functions of sRNA in mediating TE recognition and deletion (Allen and Nowacki, 2020; Couzin, 2002; Drews et al., 2022; Noto and Mochizuki, 2017). As exemplified by the characterization of heterochromatin-dependent piRNA precursor transcription mechanism in flies (Andersen et al., 2017), dissecting such unconventional transcription could gain insights into novel transcription regulatory mechanisms. However, the molecular basis by which the otherwise silent and compact MIC chromatin gains transcriptional activity in ciliates remains elusive. To date, only a limited number of studies from *Paramecium tetraurelia* and *Tetrahymena thermophila* have provided clues to understand this process. Among them, a conjugation-specific variant of the transcriptional elongation factor Spt5, complexed with Spt4, enables entry into the *Paramecium* meiotic MIC and facilitates transcription (Gruchota et al., 2017; Owsian et al., 2022). While in *Tetrahymena*, Emit1, Emit2, and Emit3 enable MIC ncRNA transcription either by promoting the localization of the transcriptional Mediator complex to the meiotic MIC (Tian et al., 2019) or by stabilizing Pol II on chromatin (Cai et al., 2025). Another Mediator complex-associated protein, Rib1, is proposed to guide the preferential Pol II-dependent transcription at TE-enriched regions (Noto et al., 2015; Schoeberl et al., 2012; Tian et al., 2019). A previous study has ruled out the involvement of trans-generational epigenetic inheritance in defining transcriptional specificity in *Tetrahymena* (Noto and Mochizuki, 2018). However, the sequential features linked to *Tetrahymena* MIC ncRNA transcription, and whether and how Rib1 promotes transcription initiation at these sites, remain unexplored.

To address the above questions, we performed a comprehensive analysis of meiotic ncRNA sequencing data, revealing that regions with consecutive G/C-rich tracts are positively associated with active transcription. Additionally, through systematic chromatin profiling and protein truncation analyses, we discovered that Rib1 promotes TE-biased ncRNA transcription by facilitating the accumulation of Pol II at G/C-rich regions, likely via phase-separation.

## Results

### The sites of meiotic scnRNA production are associated with putative G-quadruplex-forming sequences containing G/C-rich tracts

To identify potential sequence features associated with regions of abundant meiotic ncRNA transcription, we retrieved small non-coding RNA sequencing data from WT meiotic cells (Noto et al., 2015). After mapping the scnRNA reads to the MIC genome, we identified a total of 1546 regions with highly enriched scnRNA reads (hereinafter referred to as scnRNA peaks), with sizes ranging from 0.51 to 103.22 kb (median: 2.82 kb; Figure 1A). As illustrated in Figure 1B for a representative region of the MIC chromosome 2, the scnRNA peak regions, along with their summits, were precisely identified. This conclusion is further supported by the preferential accumulation of scnRNA reads at all scnRNA peaks, a pattern not observed in an equivalent number of randomly selected regions (Figure S1A).

Next, we asked whether the scnRNA peaks share any consensus DNA sequence features. To address this question, all scnRNA peak sequences were extracted and analyzed for consensus motif identification using RSAT (Santana-Garcia et al., 2022). This analysis revealed that, compared to 1546 of random MIC genomic regions, G-rich (e value = 0) and C-rich motifs (e value = 0) are significantly enriched within scnRNA peaks (Figure S1B). A further statistical analysis revealed that 76.32% of scnRNA peaks contain either a GGGGG motif (5G) or a CCCCC motif (5C, Figure S1C).

To determine whether regions containing 5G/5C tracts exhibit higher levels of meiotic scnRNA reads, we identified all MIC regions with 5G/5C tracts, then evaluated the scnRNA read coverage at and surrounding these regions. For comparison, the distribution of scnRNA reads at an equivalent number of randomly selected regions was also analyzed. Data shown in Figure 1C demonstrates that scnRNA reads are significantly enriched at sites containing 5G or 5C tracts, while no such enrichment was observed at equivalent numbers of random sites. These data imply that G/C rich tracts are prevalent features of the actively transcribed regions in the meiotic nucleus.

We noticed that scnRNA reads are markedly lower at 5G/5C tracts and their close proximity regions than their surrounding regions. This reduced sequencing read coverage at these G/C-rich sites indicates that they may be positively correlated with transcription initiation (see below). scnRNA reads showed twin peaks with a slightly higher coverage at downstream of the 5C tract and a slightly higher coverage at upstream of the 5G tract, indicating that the bidirectional ncRNA transcriptions may have a moderate preference towards regions downstream of the 5C tract.

Because there are plenty of MIC regions containing 5G/5C tracts, we sought to find out whether all 5G/5C tracts containing regions have abundant meiotic scnRNA reads. To do this, we determined scnRNA read counts within 1 kb regions surrounding all 5G/5C tracts containing sites and equivalent numbers of random sites. As illustrated in Figure 1D, 69.66% of 5G or 69.72% of 5C motif-containing regions have more than 10 scnRNA read counts. Although these percentages are radically lower than those of random sites (around 30%), the results still imply that, although the presence of 5G/5C tracts is highly correlated with the scnRNA production, additional features other than G/C-rich motifs jointly determine the active transcription regions in the MIC.

Given that G/C-rich DNA is prone to form G-quadruplex, a four-stranded secondary DNA structure (Lombardi and Londono-Vallejo, 2020; Spiegel et al., 2020), and ample recent evidence has shown that G-quadruplex promotes transcription (Esain-Garcia et al., 2024; Wang et al., 2024), we then identified potential G-quadruplex-forming sequences (PQSs) in the MIC genome and investigated whether regions containing both 5G/5C tracts and PQSs exhibit a stronger positive correlation with active ncRNA transcription. The results showed that the level of scnRNA reads enrichment at PQSs is comparable with that of regions with 5G/5C tracts (Figure 1C-D). However, scnRNA reads have a markedly higher enrichment at PQSs that contain 5G/5C tracts (Figure 1C-D).

Having found 5G/5C tracts containing PQSs (hereinafter referred to as 5G/5C-PQSs) exhibited a stronger positive correlation with active MIC ncRNA transcription, we then studied their spatial relationships with IESs. To achieve this, we focused on two groups of IESs: Type A1 IESs, which produce abundant scnRNAs during the early stage of conjugation (i.e., the meiotic prophase, primarily derived from the MIC), and Type B2 IESs, which predominantly produce scnRNAs in the late stage of conjugation, during the formation of new MACs ((Noto et al., 2015), Figure 1E). They are selected for the analysis primarily due to their distinct temporal patterns of transcription, which would allow us to figure out whether 5G/5C-PQSs are specific to the early expressed IESs in meiosis. Besides, their numbers are comparable: There are 1899 Type A1 and 1654 Type B2 IESs, respectively. After analyzing the distribution of 5G/5C-PQSs at and around two Types of IESs, we found that they showed a strong enrichment at the termini of Type A1 IESs, along with moderate enrichment in the middle of Type A1 IESs (Figure 1F and Figure S1D). By contrast, a similar enrichment was not observed at Type B2 IESs (Figure 1F). Summing up, 5G/5C-PQSs likely serve as cis-elements that promote the MIC ncRNA transcription in a subset of Type A1 IESs (Figure 1G).

### RNA polymerase II is preferentially associated with G/C-rich motifs in the meiotic MIC

To find out whether the Pol II transcriptional machinery is preferentially associated with 5G/5C-PQSs, we sought to characterize Pol II binding sites in the meiotic MIC, using Cleavage Under Targets & Tagmentation (CUT&Tag, (Kaya-Okur et al., 2019)). Briefly, we prepared meiotic cells expressing Rpb3 with a C-terminal HA tag under control of its endogenous promoter. Immunostaining revealed that Rpb3 (by an anti-HA antibody) and double-stranded RNA (dsRNA; i.e., scnRNA precursors; labelled by an anti-dsRNA antibody) shared a largely overlapping localization pattern in the meiotic MIC (Figure 2A-B), indicating that Rpb3 is enriched at regions with ncRNA biogenesis.

**Figure 2.**
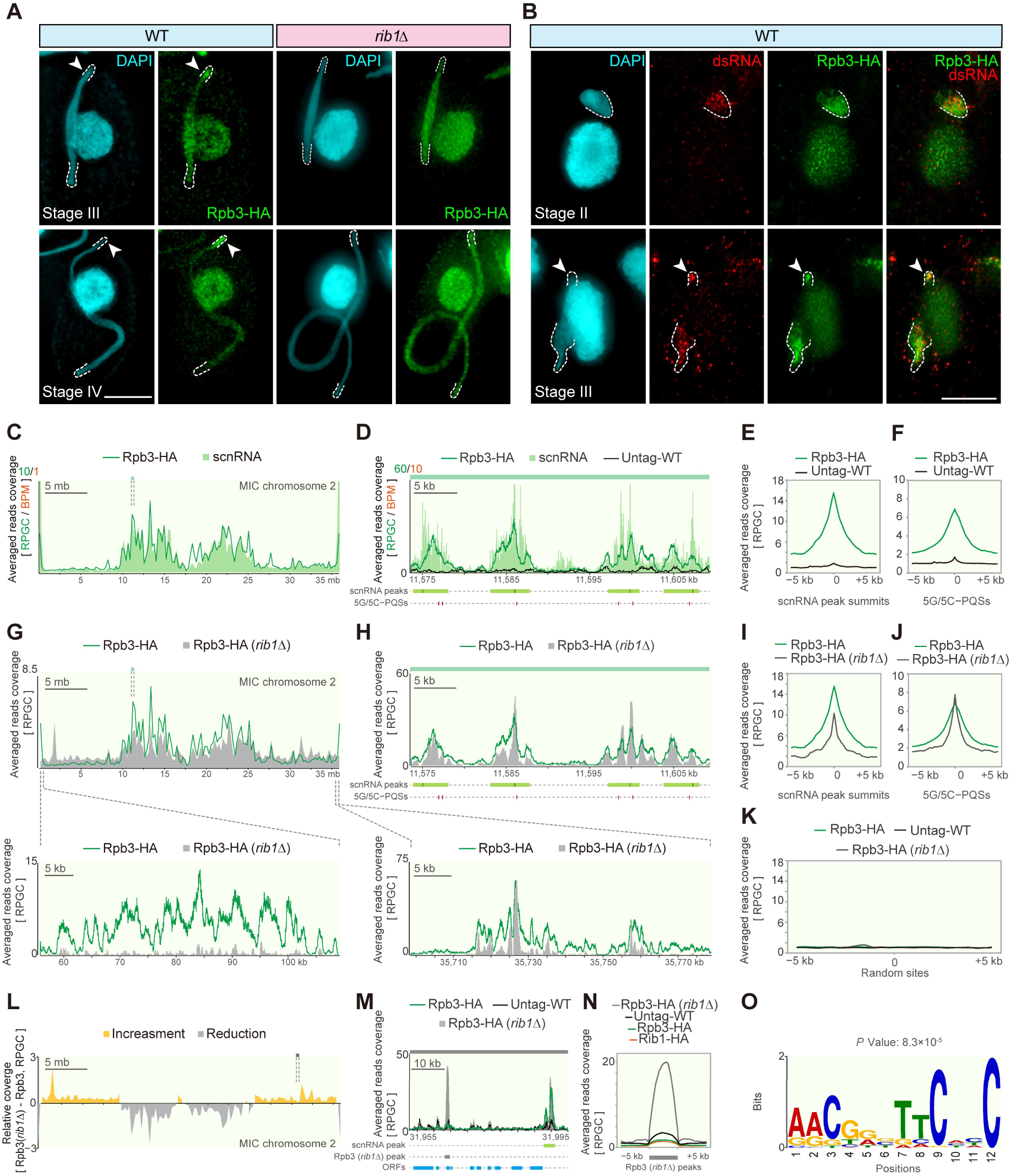
RNA polymerase II binds scnRNA production regions and such specificity are partially dependent on Rib1. A, and B, Immunostaining of the endogenously expressed Rpb3-HA fusion protein revealed its prominent co-localization with double-stranded RNAs (dsRNA) in the meiotic MIC. This localization is disrupted by *RIB1* deletion. Dashed lines indicate the MIC termini in panel A and the dsRNA-enriched regions in panel B. Arrowheads mark regions with enriched Rpb3-HA. Scale bars: 10 μm. C, Normalized Rpb3-HA CUT&Tag read coverage and the WT meiotic scnRNA sequencing read coverage on MIC chromosome 2. CUT&Tag and scnRNA sequencing reads were normalized using ‘reads per genomic content (RPCG)’ and ‘bins per million mapped reads (BPM)’, respectively. The green box denotes the region selected for magnification in panel D. D, Normalized Rpb3-HA CUT&Tag read coverage and WT meiotic scnRNA sequencing read coverage for a region of MIC chromosome 2 (coordinate: CM028432.1:11,599,915−11,617,719). Green bars represent meiotic scnRNA peaks, and black vertical lines indicate peak summits. Red vertical lines mark sites containing 5G/5C-PQSs. E, Visualization of averaged CUT&Tag read coverage at and surrounding the meiotic scnRNA peak summits. F, Visualization of averaged CUT&Tag read coverage at and surrounding 5G/5C-PQSs. G, Upper panel: Normalized Rpb3-HA CUT&Tag read coverage in WT or *rib1*Δ MIC chromosome 2. Lower panels: Magnified visualizations of CUT&Tag read coverages at the termini of MIC chromosome 2. The green box denotes the region selected for magnification in panel H. H, Normalized Rpb3-HA CUT&Tag read coverage and WT meiotic scnRNA sequencing read coverage for a region of MIC chromosome 2. I, Visualization of averaged CUT&Tag read coverage at and surrounding the meiotic scnRNA peak summits. J, Visualization of averaged CUT&Tag read coverage at and surrounding 5G/5C-PQSs. K, Visualization of averaged CUT&Tag read coverage at and surrounding random sites. L, Relative CUT&Tag read coverage showing the increase and decrease of Rpb3 occupancy on MIC chromosome 2 in response to *RIB1* deletion. The grey box highlights the region selected for magnification in panel M. M, Normalized CUT&Tag read coverage in a region with aberrant Rpb3 occupancy in *rib1*Δ cells (coordinate: CM028432.1:31,456,953−31,469,015). Gray bars represent aberrant Rpb3 peaks, and black vertical lines indicate peak summits. N, Visualization of averaged CUT&Tag read coverage at and surrounding the aberrant Rpb3 peaks found in *rib1*Δ cells. O, A DNA motif identified from aberrant Rpb3 peaks.

Next, we purified MICs from Rpb3-HA meiotic cells and used these as starting materials for CUT&Tag (Figure S2A). As a control, a mock CUT&Tag was performed using MICs purified from untagged WT meiotic cells. Figure 2C illustrates the coverage of Pol II CUT&Tag reads at MIC chromosome 2. Pol II exhibits a scnRNA-like pattern across the entire chromosome. Locally, Pol II preferentially occupies the scnRNA peaks (Figure 2D). In contrast, a mock CUT&Tag performed with the same amount of anti-HA antibody on MICs purified from untagged WT meiotic cells showed no preferential accumulation at the scnRNA peaks (Figure 2D-E). Pol II CUT&Tag read coverage profile at and surrounding all scnRNA peak summits further revealed that the preferential accumulation of Pol II at the meiotic scnRNA production sites is a general feature across the genome (Figure 2E). This observation is consistent with the largely overlapping localizations of Rpb3 and dsRNAs in the meiotic MICs (Figure 2B).

We then investigated Rpb3 occupancy at 5G/5C-PQSs. Like the meiotic scnRNAs, Rpb3 exhibits enriched occupancy at these sites, but unlike the meiotic scnRNAs, the reduction of occupancy at the center of 5G/5C-PQSs was not detected for Rpb3 (Figure 2F). Since Pol II typically accumulates near transcription start sites, these findings suggest that 5G/5C-PQSs may positively correlate with transcription initiation in the MIC.

### Rib1 assures the specific accumulation of RNA polymerase II at the meiotic scnRNA peak regions

Although the above analyses identified cis-elements positively correlated with the MIC ncRNA transcription, the transcriptional regulator that binds to these sites and recruits Pol II for transcription initiation remains unknown. During MIC transcription, the chromosomes undergo drastic rearrangement, with telomeres and centromeres clustered at opposite poles of the elongated nucleus (Loidl et al., 2012; Tian et al., 2022). Consequently, IESs, which are highly enriched in subtelomeric and pericentromeric regions, exhibit a nonhomogeneous localization pattern in the MIC. They occupy a narrow region at one pole of the elongated MIC while being broadly distributed at the opposite pole (Tian et al., 2019). Our previous work suggested that Rib1, a Mediator complex-associated protein, is essential for the preferential accumulation of Pol II at opposite poles of the elongated meiotic MIC, where IESs are highly enriched, and important for the MIC ncRNA biogenesis (Hamilton et al., 2016; Tian et al., 2019) (Figure 2A). These observations prompted us to investigate whether Rib1 plays a role in recruiting Pol II to the cis-elements for the scnRNA production.

Consistent with our cytological observations, CUT&Tag analysis showed that Rpb3 was less enriched at the scnRNA production regions in *rib1*Δ cells (Figure 2G and Figure S2B-S2E). However, Rib1 deletion did not abolish Rpb3 occupancy at scnRNA peaks (Figure 2H). Further comparison of averaged Rpb3 occupancy at all scnRNA peak summits (Figure 2I) and at all 5G/5C-PQSs revealed that Rpb3-occupied regions became narrower on *rib1*Δ MIC chromatin (Figure 2J-K). These findings suggest that Rib1 is not essential for the association of Pol II with the majority of scnRNA production sites but may be required for the efficient Pol II elongation.

Another notable change in Rpb3 occupancy following Rib1 deletion was its aberrant accumulation at scnRNA-poor regions (Figure 2L), along with a partial reduction - but not complete loss - of Pol II occupancy at chromosome ends in *rib1*Δ cells (Figure S2F). Notably, Rpb3 was not uniformly distributed across these regions but instead showed enrichment at specific loci (Figure 2M). We systematically identified such representative aberrant Pol II binding regions through a comparative analysis of Rpb3 occupancy in WT and *rib1*Δ MIC chromatin. This analysis identified 1,241 confident regions (Figure 2M). CUT&Tag read coverage comparison confirmed that Rpb3 specifically associates with these regions in *rib1*Δ cells, whereas Rpb3 does not accumulate at these regions in WT cells (Figure 2N). Although motif analysis using the STREAM algorithm (Bailey, 2021) retrieved a motif (Figure 2O, *p*-value = 8.3 × 10⁻⁵), it present in only 32.72% of the analyzed regions. Therefore, the abnormal accumulation of Rpb3 at specific scnRNA-poor regions in *rib1*Δ meiotic chromatin is unlikely to be driven solely by DNA sequence features. Altogether, we conclude that Rib1 is not required for Pol II binding to meiotic scnRNA peaks, but is essential for limiting the Pol II association to these loci.

### Rib1 preferentially occupies the MIC ncRNA transcription region

Given that the disruption of Rib1 resulted in aberrant accumulation of Rpb3 at regions outside of the meiotic scnRNA production sites, and considering Rib1’s co-localization with Rpb3 (Figure 3A), we hypothesized that Rib1 is required for promoting the accumulation of Pol II at these transcription sites. To test this idea, we examined Rib1 chromatin localization. We purified MICs from meiotic cells expressing endogenous levels of C-terminally HA-tagged Rib1 and performed CUT&Tag analysis for Rib1-HA. While Rib1 did not show obvious enrichment at the non-transcribed regions where Rpb3 aberrantly accumulated in *rib1*Δ (Figure 2N), it preferentially occupied regions enriched in Rpb3 (Figure 3B-C), which is consistent with the cytological co-localization of these proteins in the MIC (Figure 3A). CUT&Tag read coverage profile analysis further showed overlapping occupancy of Rib1 and Rpb3 at scnRNA peaks (Figure 3D) and at the 5G/5C-PQS sites in the WT MIC chromatin (Figure 3E) but not at an equivalent number of random sites (Figure 3F). These findings suggest that Rib1 is specifically enriched at scnRNA peaks to promote the specific association of Pol II at the meiotic scnRNA production sites, and consequently reduces Pol II’s non-specific chromatin binding indirectly.

**Figure 3.**
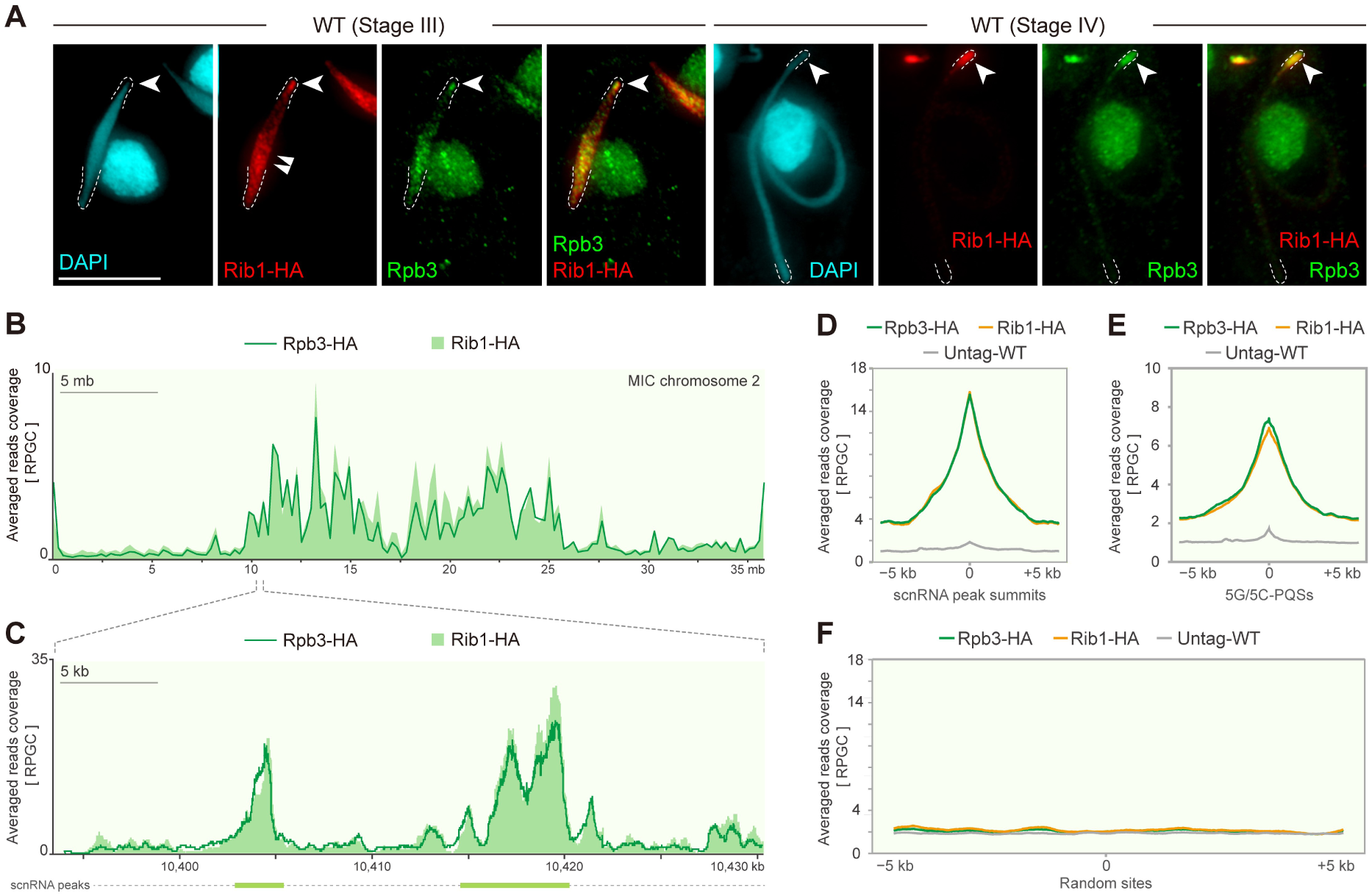
Rib1 preferentially associates with the regions involved in the production of meiotic scnRNA. A, Co-immunostaining of endogenously expressed Rib1-HA with Rpb3. Dashed lines indicate the MIC termini. The single arrowheads point to accumulated Rib1-HA and Rpb3 at a narrow region of the MIC, while the double arrowheads denote accumulated Rib1 at a broader region of the MIC. Scale bar: 10 μm. B, Normalized Rpb3-HA and Rib1-HA CUT&Tag read coverage on MIC chromosome 2 in wild-type (WT) cells. CUT&Tag reads were normalized using ‘reads per genomic content (RPCG)’. The blue box denotes the region selected for magnification in panel C. C, Normalized CUT&Tag read coverage for a region of MIC chromosome 2. Light blue bars represent meiotic scnRNA peaks. D, Visualization of averaged CUT&Tag read coverage at and surrounding the meiotic scnRNA peak summits. E, Visualization of averaged CUT&Tag read coverage at and surrounding 5G/5C-PQSs. F, Visualization of averaged CUT&Tag read coverage at and surrounding random sites.

### Rib1’s disordered N-terminal and folded C-terminal domains coordinate meiotic MIC localization and scnRNA-dependent DNA elimination

To find out how Rib1 promotes the preferential accumulation of Pol II, we analyzed its protein sequence. Rib1 is a 424 amino acid protein with no identifiable protein domain. Sequence analyses using PLAAC and IUPred3 suggest that the N-terminal domain (NTD, 1–272 aa) is intrinsically disordered and has a high potential to form a prion-like structure (Figure 4A), a structure that may drive the formation of protein condensates (Alberti et al., 2018; Boeynaems et al., 2018). Moreover, three phase-separation-driving peptides within NTD were identified using PSPhunter (Figure 4A, (Sun et al., 2024)). Besides, the NTD contains nine tandemly arranged N/Q-rich motifs [NQMNQN]. While similar repeats have been identified in Mediator complex subunits from other organisms (Tian et al., 2019), their biological function remains unclear. In contrast to the unstructured N-terminal region, the AlphaFold prediction suggests that the C-terminal domain is folded and consists of multiple alpha helices (Figure S3A, (Jumper et al., 2021; Varadi et al., 2024)).

**Figure 4.**
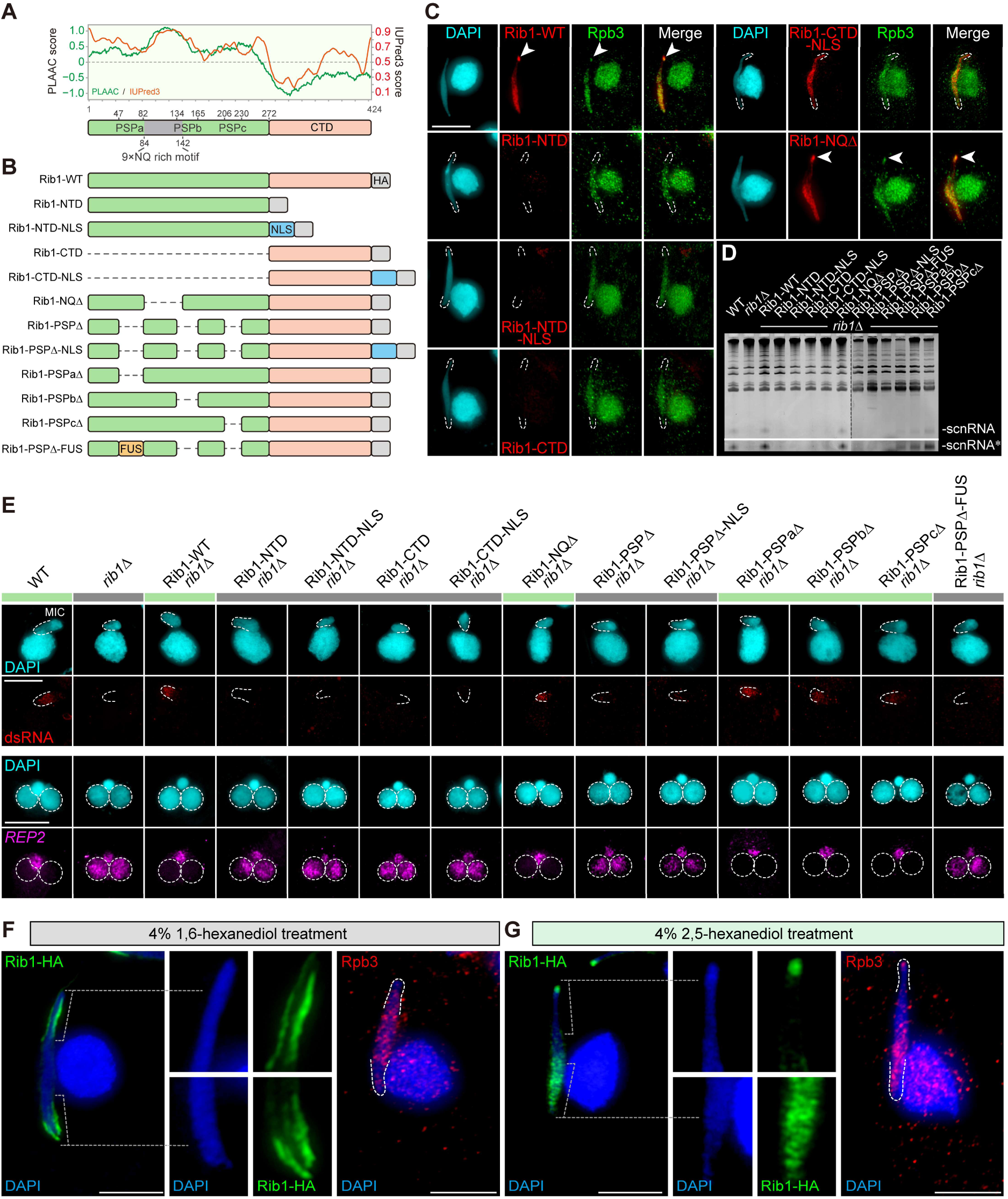
Truncation analysis of Rib1 protein. A, Upper panel: Putative prion-like sequences and intrinsically disordered regions predicted by PLAAC and IUPred3, respectively. Lower panel: Schematic representation of the protein domains, including the disordered N-terminal domain (NTD, green), the folded C-terminal domain (CTD, orange), putative phase-separating peptides identified using PSPhunter, and the NQ-rich motif-containing region (grey). B, Schematic diagrams of Rib1 variants generated in this study. HA indicates the hemagglutinin epitope tag. NLS represents a characterized nuclear localization signal peptide specific to *Tetrahymena* MIC. FUS denotes the phase-separating peptide derived from the human FUS protein. C, Co-Immunostaining of HA-tagged Rib1 variants and Rpb3. Arrowheads indicate positions with accumulated Rib1 (or Rib1 variants) and Rpb3. Scale bar: 10 μm. D, Analysis of scnRNA biogenesis by RNA gel electrophoresis. Total RNAs were all isolated from meiotic cells. The asterisk denotes longer exposure. Uncropped images are listed in Figure S4B. E, Analysis of dsRNA biogenesis and DNA elimination in WT, *rib1*Δ, and *rib1*Δ cells expressing different Rib1 variants. DsRNA biogenesis was assessed by immunostaining. DNA elimination was monitored by fluorescence *in situ* hybridization (FISH). Dashed lines indicate DAPI-poor regions of the slightly elongated meiotic MICs, while dashed circles mark the new MACs. Green bars indicate dsRNAs are detectable and *REP2* IESs were removed from the new MACs. While grey bars indicate dsRNA biogenesis and *REP2* IESs elimination are defective. F, and G, Immunostaining of Rib1-HA and Rpb3 in cells treated with 1,6-hexanediol or 2,5-hexanediol. Scale bars: 10 μm.

To further investigate how the different regions of Rib1 contribute to its localization and function, we generated a series of *rib1*Δ cells expressing truncated Rib1 variants (Figure 4B and Figure S3B-C). In parallel, *rib1*Δ cells expressing wild-type Rib1 (Rib1-WT) were also generated as controls. Successful protein expression was confirmed by Western Blotting (Figure S3D).

We first examined whether the truncations affect the Rib1 localization by indirect immunostaining of the HA-tagged Rib1 variants. We found that truncations of either the NTD or CTD prevented Rib1 from localizing to the meiotic MIC. We then generated additional strains expressing either Rib1-NTD or Rib1-CTD fused with a characterized micronuclear localization signal (MicNLS2, (Iwamoto et al., 2018)) and found that the MicNLS2 sequence can restore the MIC localization of Rib1-CTD, but not Rib1-NTD (Figure 4C).

We next examined Rpb3 localization in *rib1*Δ cells expressing the different Rib1 variants. Rpb3 displayed a WT-like, nonhomogeneous pattern in Rib1-WT cells but an abnormal, homogeneous pattern in Rib1-NTD, Rib1-NTD-NLS, and Rib1-CTD cells (Figure 4C). Correspondingly, dsRNA was detected in Rib1-WT meiotic MICs but absent in Rib1-NTD, Rib1-NTD-NLS, and Rib1-CTD cells, leading to undetectable scnRNA (Figure 4D-E). Consequently, although DNA elimination of *REP2*-complementary IESs was completed in the progeny of Rib1-WT cells, they were retained in Rib1-NTD, Rib1-NTD-NLS, and Rib1-CTD cells (Figure 4E). These results indicate that meiotic scnRNA biogenesis and DNA elimination are abolished in these mutants, consistent with the impaired Rib1 localization to the meiotic MIC.

We tested whether Rib1-CTD-NLS (which localizes to the MIC) could rescue the MIC ncRNA transcription defect in *rib1*Δ cells. Despite MIC localization, Rib1-CTD-NLS showed abnormal homogeneous distribution (Figure 4C) and failed to restore Rpb3’s nonhomogeneous patterning, dsRNA/scnRNA biogenesis (Figure 4D-E), and *REP2* IES elimination in new MACs (Figure 4E). These results demonstrate that both the disordered N-terminal and structured C-terminal domains are essential for Rib1-mediated meiotic ncRNA production.

### The intrinsically disordered region of Rib1 is essential for its function in transcriptional regulation

Given that Rib1 shows preferential aggregation at the peri-centromeric regions and telomeric pole (Figure 3A), we were prompted to investigate whether the Rib1 aggregates are sensitive to 1,6-hexanediol treatment, a reagent that is commonly used to disrupt liquid condensates formed via phase separation (Kroschwald et al., 2017; Liu et al., 2021). To assess this, we treated Rib1-HA cells with 4% 1,6-hexanediol. For comparison, cells from the same batch were also treated with 4% 2,5-hexanediol, a similar aliphatic alcohol that is less effective in disrupting liquid condensates (Guzikowski and Kavalali, 2024; Lin et al., 2016).

Immunostaining results revealed that 1,6-hexanediol treatment severely disrupted Rib1 aggregation on chromatin (Figure 4F) and caused abnormal localization of Rib1 to the periphery of the MIC. In line with the abolished Rib1 foci on the MIC chromatin, Rpb3 showed an abnormal, homogeneous distribution within the MIC. Unlike 1,6-hexanediol-treated cells, such aberrant localization patterns of Rib1 or Rpb3 were not observed in cells treated with 2,5-hexanediol (Figure 4G). Altogether, the sensitivity of Rib1 foci to 1,6-hexanediol treatment suggests that Rib1’s localization in the meiotic MIC may be regulated through phase separation.

Next, we dissected which subdomain(s) of the disordered Rib1 NTD are essential for its function. We introduced Rib1-NQΔ, which lacks all NQ-rich motifs (Figure 4B), into *rib1*Δ cells and analyzed its function. Rib1-NQΔ was capable of localizing to the MIC and exhibited a WT-like, nonhomogeneous pattern in the meiotic MIC (Figure 4C). Rib1-NQΔ restored Rpb3’s nonhomogeneous localization, dsRNA and scnRNA biogenesis in *rib1*Δ meiotic MICs (Figure 4D-E) and showed elimination of *REP2* IESs in new MACs (Figure 4E).

The successful rescue of *rib1*Δ deficiency by the NQ-rich motif-deleted version of Rib1 prompted us to investigate whether the absence of the NQ-rich motif might lead to subtle defects in Rib1 chromatin occupancy, which might not be detectable by immunostaining. To address this, we isolated meiotic MICs expressing Rib1-WT and Rib1-NQΔ and used these for CUT&Tag analysis. Additionally, MICs from meiotic cells expressing Rib1-CTD-NLS were harvested as a negative control for the CUT&Tag experiment. As shown in Figure 5A-D, deletion of the NTD (i.e., in Rib1-CTD-NLS cells) severely impaired Rib1’s association with meiotic chromatin. In contrast, Rib1-WT and Rib1-NQΔ exhibited highly overlapping chromatin occupancy patterns, both locally and globally, suggesting that the NQ-rich repeats are dispensable for mediating Rib1’s association with meiotic chromatin.

**Figure 5.**
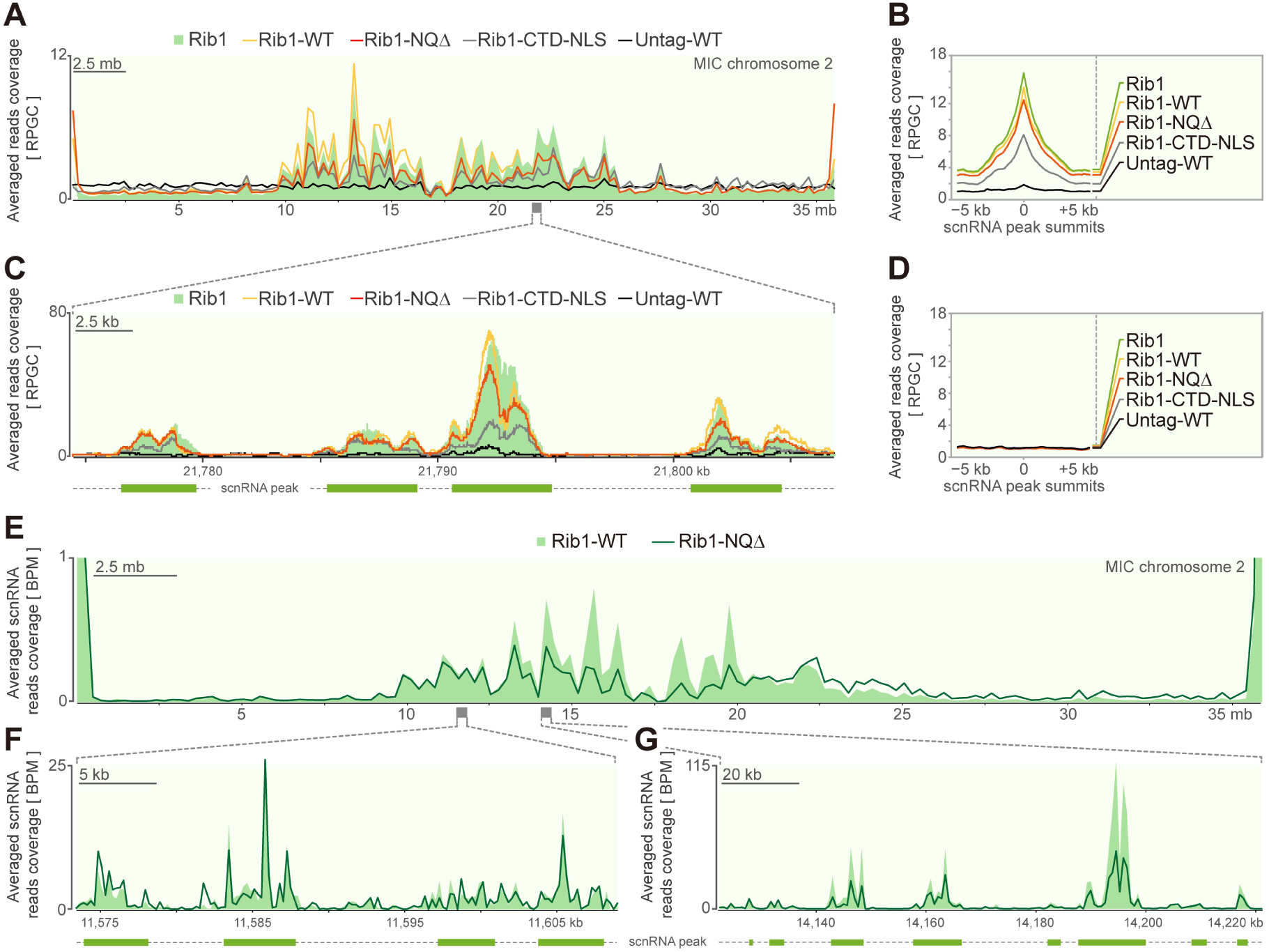
Chromatin profiling analysis of Rib1 and its variants. A, Normalized CUT&Tag read coverage on MIC chromosome 2. CUT&Tag reads were normalized using ‘reads per genomic content (RPCG)’. The grey box denotes the region selected for magnification in panel C. B, Visualization of averaged CUT&Tag read coverage at and surrounding the meiotic scnRNA peak summits. C, Normalized CUT&Tag read coverage for a region of MIC chromosome 2 (coordinate: CM028432.1:21,774,510−21,806,788). Green bars represent meiotic scnRNA peaks. D, Visualization of averaged CUT&Tag read coverage at and surrounding random sites of the MIC genome. E, Normalized scnRNA sequencing read coverages on the MIC chromosome 2. The scnRNA sequencing reads were normalized using ‘bins per million mapped reads (BPM)’. Grey boxes denote the regions selected for magnification in panel F and G, respectively. F,G, Normalized scnRNA sequencing read coverage at two representative regions of MIC chromosome 2. Green bars indicate regions corresponding to meiotic scnRNA peaks. Their coordinates are CM028432.1:11,573,403−11,609,011 (panel F) and CM028432.1:14,122,517−14,221,025 (panel G).

Next, we sought to determine whether Rib1-NQΔ could restore scnRNA biogenesis at the correct loci and in appropriate amounts. To do so, we extracted total RNA from Rib1-WT and Rib1-NQΔ strains and performed small RNA sequencing analysis. The results indicated that both strains expressed comparable levels of small RNAs from consensus regions (Figure 5E-F). However, the deletion of the NQ-rich motif did impair scnRNA biogenesis at certain regions, especially at regions close to the middle of the chromosomes (Figure 5E and Figure 5G). Taken together, these findings suggest that, while Rib1 contains evolutionarily conserved NQ-rich repeats, these repeats are not essential for Rib1’s chromatin occupancy or for Rib1-directed small RNA biogenesis.

Because the NQ-rich region constitutes only a portion of Rib1’s intrinsically disordered region (IDR) and is nearly mutually exclusive with the putative phase-separation-driving peptides (PSPs), the above data do not exclude the possibility that PSPs are essential for Rib1’s function.

Next, we asked whether deletion of PSPs impairs Rib1’s function. To this end, we generated *rib1*Δ cells expressing Rib1 variant with all PSPs deleted (Rib1-PSPΔ). The immunostaining result showed that Rib1-PSPΔ was not detected in the meiotic MIC (Figure 6A), suggesting that PSPs are essential for the localization of Rib1 to the meiotic MIC. Thus, to study the function of Rib1-PSPs, we generated Rib1-PSPΔ with a MIC localization signal (Rib1-PSPΔ-NLS). Immunostaining data showed that the addition of the MicNLS2 to Rib1-PSPΔ tethered the protein to the meiotic MIC (Figure 6B). Nevertheless, it exhibited an abnormal homogeneous pattern in the MIC. Similarly, Rpb3 showed an aberrant localization pattern in the MIC. To further narrow down which PSP is essential for Rib1 to form the nuclear aggregation, we generated three more *rib1*Δ cells expressing Rib1 protein missing one of the PSPs (i.e., Rib1-PSPaΔ, Rib1-PSPbΔ, and Rib1-PSPcΔ), then investigated their localizations. The results revealed that removing the PSPa did not notably affect the localizations of Rib1 and Rpb3 (Figure 6C), while deleting PSPb or PSPc led to a slightly aberrant homogeneous localization of Rib1. Correspondingly, Rpb3 became more homogeneous in both Rib1-PSPbΔ and Rib1-PSPcΔ cells (Figure 6D-E), especially in Rib1-PSPcΔ cells. A further examination of scnRNA, dsRNA biogenesis, and *REP2* DNA elimination in the above mutants showed that dsRNA was barely detected in Rib1-PSPΔ and Rib1-PSPΔ-NLS cells. Consequently, scnRNA was not detected in these two mutants, and *REP2* IESs were abnormally retained (Figure 4D-E). By contrast, truncated Rib1 that was missing any one of the three PSPs was capable of restoring *rib1*Δ defects in the MIC ncRNA biogenesis and *REP2* IES elimination (Figure 4D-E). Taken together, while missing all PSPs severely impedes Rib1’s function, retaining any one of the two PSPs is enough for Rib1 to mediate the small RNA mediated DNA elimination pathway in *Tetrahymena*.

**Figure 6.**
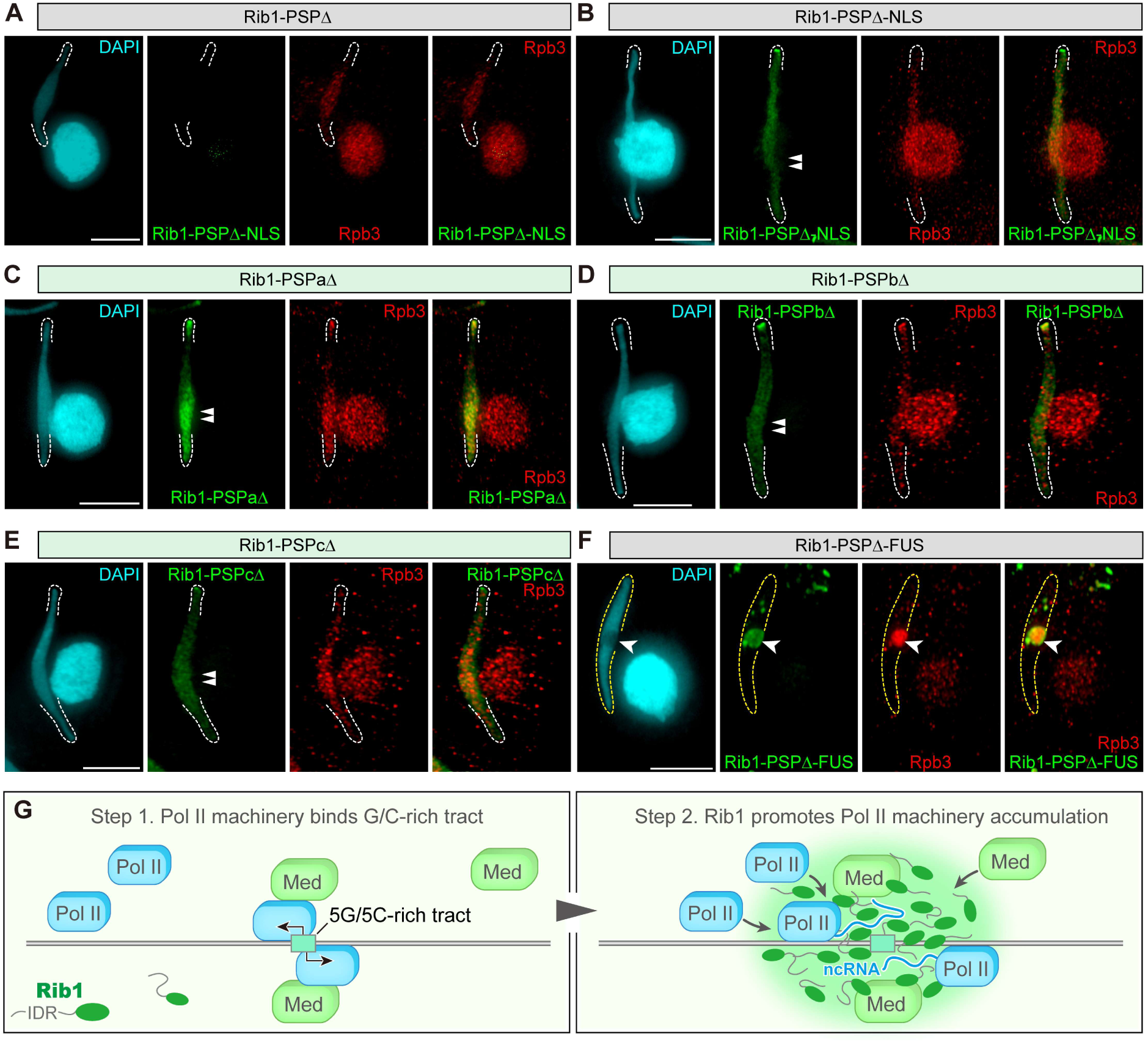
Rib1 may facilitate the accumulation of Rpb3 at meiotic MIC chromatin regions through phase separation. A,B,C,D,E, and F, Co-immunostaining of HA-tagged Rib1 variants and Rpb3 in *rib1*Δ cells. Dashed lines indicate the termini of the meiotic MIC. Double arrowheads indicate regions in the middle of the elongated MICs that are supposed to have abundant Rib1 in the WT (see Figure 3A). Single arrowheads denote Rib1 and Rpb3 aggregates in the MIC. Scale bars: 10 μm. G, Schematic diagram illustrating Rib1-mediated accumulation of the Pol II machinery at meiotic scnRNA production regions. ‘IDR’: Intrinsically disordered region, ‘ncRNA’: Non-coding RNA, ‘Pol II’: RNA polymerase II, and ‘Med’: Mediator complex.

To test whether Rib1-PSPΔ’s aberrant localization and defective function in promoting the meiotic ncRNA transcription is solely due to its loss of the aggregation ability, we generated *rib1*Δ strains expressing Rib1-PSPΔ fused to a human FUS-derived peptide known to drive phase separation (Rib1-PSPΔ-FUS, Figure 4A; (Murthy et al., 2019)). The results showed that Rib1-PSPΔ-FUS localizes to the MIC without the MicNLS2 and forms large abnormal aggregates in the MIC (Figure 6F). This suggests that the failure of Rib1-PSPΔ to localize in the MIC is unlikely due to a defect in nuclear import, but its aggregation capability is likely essential for retaining Rib1 within the MIC. Strikingly, Rpb3 was also enriched in the Rib1-PSPΔ-FUS aggregates. By contrast, a similar aggregation of Rpb3 was not observed in *rib1*Δ cells expressing the FUS-EGFP fusion protein, suggesting that both FUS and Rib1 are required for Rpb3 aggregation in the MIC (Figure S4A). Nevertheless, cytological experiments showed that Rib1-PSPΔ-FUS is not functional, as neither MIC ncRNA transcription nor the *REP2* IES elimination was restored (Figure 4D-E). In summary, tightly regulated aggregation of Rib1 may be crucial for its role in facilitating proper Pol II localization, which in turn mediates meiotic ncRNA transcription in the MIC (Figure 6G).

## Discussion

### Potential G-quadruplex-forming sequence with G/C-rich tracts exhibits a strong positive correlation with the MIC ncRNA transcription

Transcription of the *Tetrahymena* meiotic MIC is highly biased towards recent integrated IESs (Noto et al., 2015), yet the mechanism underlying this preference was unclear. Through the identification of MIC regions enriched with meiotic scnRNA reads and subsequent DNA sequence feature analyses, we discovered that 5G/5C-PQSs exhibit a strong positive correlation with the meiotic ncRNA transcription (Figure 1). Correspondingly, Pol II preferentially associates with regions exhibiting these sequence features (Figure 2F).

What is the role of 5G/5C-PQSs in the MIC ncRNA transcription? Although G-quadruplexes have long been considered as transcriptional ‘roadblocks’, growing evidence demonstrates that they can promote transcription both *in vivo* and *in vitro* (see (Robinson et al., 2021) and (Chen et al., 2024; Lee et al., 2020)). First, G-quadruplexes are highly enriched at nucleosome-depleted promoter regions of actively transcribed genes in humans (Hansel-Hertsch et al., 2016; Li et al., 2021a), likely because they facilitate nucleosome exclusion (Esain-Garcia et al., 2024). Second, G-quadruplexes serve as binding platforms for transcription factors, which in turn facilitate transcription (Lago et al., 2021; Li et al., 2021b; Spiegel et al., 2021; Wulfridge et al., 2023; Yuan et al., 2023). Third, transcripts originating from G-quadruplex-containing regions are prone to forming stable R-loops, which may help keep the DNA double helix open and facilitate transcription reinitiation (see (Lee et al., 2020; Wulfridge and Sarma, 2024)). Moreover, recent studies suggest that R-loops may play a role in regulating chromatin dynamics (Al-Hadid and Yang, 2016; Liu et al., 2024; Liu et al., 2020). Therefore, similar mechanisms may explain the preferential transcription at 5G/5C-PQSs in *Tetrahymena* MIC.

Why do scnRNA reads show low coverage at the center of 5G/5C-PQSs if they facilitate the MIC ncRNA transcription? This observation could be explained by the following non-mutually exclusive possibilities: i) scnRNA carrying a 5’-triphosphate or 5’-cap from de novo initiation site cannot be ligated with sequencing adapter using T4 RNA ligase, so small RNA sequencing of scnRNAs does not recapitulate the transcript termini of the precursor MIC ncRNA transcripts. ii) the low coverage of scnRNA reads at 5G/5C-PQSs suggests that these sites could be transcription initiation sites where the MIC ncRNA transcripts barely form dsRNAs. iii) the post-transcriptional processing and loading mechanism of scnRNAs makes most scnRNAs have uracil as the first base (5′ U) (Lee and Collins, 2006; Mochizuki and Kurth, 2013). Therefore, the lower abundance of scnRNA reads at G/C-rich regions may be due to the inefficient production and loading of scnRNAs from G/C-rich dsRNAs.

What drives the adoption of such a G/C-rich motif as a signature for active scnRNA transcription? Notably, the resemblance of the 5G/5C-tracts to the MAC *Tetrahymena* telomeric repeat (T2G4) suggests a potential origin from telomeres. In *Tetrahymena*, TEs remain transcriptionally silenced in the MIC, except during meiotic prophase. Even when the MIC gains transcriptional activity during meiosis (Cai et al., 2025; Mochizuki and Gorovsky, 2004; Sugai and Hiwatashi, 1974; Tian et al., 2019), essential factors for mRNA processing and maturation are absent from the meiotic MIC (Zhao et al., 2019). Thus, the only opportunity for TEs to produce mRNA occurs after the formation of the MAC anlagen (Zhao et al., 2019). Soon after the formation of the MAC anlagen, DNA elimination factors excise over 12,000 IESs from the MAC anlagen (Hamilton et al., 2016). Concomitantly with IES elimination, the five zygotic chromosomes are fragmented into a total of 181 chromosomes (Eisen et al., 2006; Sheng et al., 2020; Wang et al., 2021; Yao et al., 1990), followed by de novo telomere addition at the broken chromosome ends (see review (Coyne et al., 1996)). The simultaneous excision of 12,000 IESs and fragmentation of hundreds of chromosomal DNA segments present a significant challenge for the telomere addition machinery. Moreover, as excised IESs are rarely cyclized in *Tetrahymena* (Saveliev and Cox, 1994, 2001), *de novo* telomere addition to some IES termini may take place. The addition of telomeres may stabilize these IESs. As transposons in IESs are mostly DNA transposons and their remnants (Hamilton et al., 2016), and IES-derived mRNA may lead to the production of transposase, consequently, it would facilitate the reintegration of telomere bearing IESs into the new MIC genome. Over time, these intrachromosomal telomeric repeats may degenerate due to higher mutation rates of repetitive sequences (e.g., from non-allelic recombination), forming G/C-rich segments at the IES termini. These G/C-rich tracts could then serve as a signature motif for promoting highly preferential transcription initiation of ncRNAs from recently integrated IESs (Noto et al., 2015).

The biological significance of G/C-rich tracts and G-quadruplex DNA in *Tetrahymena* was largely unknown. Although not directly linked to transcriptional regulation in the MIC, previous studies have shown that some IESs are flanked by AAAAAGGGGG (A_5_G_5_) sequences, and disruption of these sequences severely impairs IES elimination, at least at the tested locus (Godiska et al., 1993; Godiska and Yao, 1990). Additionally, disruption of a G-quadruplex binding protein Lia3 leads to imprecise excision of a subset of A_5_G_5_-flanked IESs in the new MAC (Carle et al., 2016). Together, these findings—along with our results—suggest that G/C-rich tracts and G-quadruplex structures serve as signature elements for IES transcription (in the parental MIC) and elimination (in the new MAC). The discovery of Lia3 also raises the possibility that similar G-quadruplex-binding protein(s) may be present in the meiotic MIC, where they could direct Pol II to 5G/5C-PQSs and initiate ncRNA transcription from IES-rich regions.

### The intrinsically disordered region of Rib1 is essential for maintaining the accumulation of Pol II at the scnRNA production sites

Our previous study proposed that Rib1 may recognize transcription activation signals and direct the preferential association of transcription machinery with IES-rich chromatin (Tian et al., 2019). In the present study, we confirmed that Rib1 is preferentially associated with the meiotic scnRNA production sites (Figure 5A). Surprisingly, however, the association of Pol II with scnRNA production sites was only moderately impaired by *RIB1* deletion (Figure 2G-J). Nevertheless, Rib1 deletion led to abnormal Pol II association with regions outside the scnRNA production sites (Figure 2L-M). Notably, Rib1 does not associate with these aberrant Pol II-binding regions (Figure 2N), which implies that Rib1 passively prevents Pol II from binding to these non-specific regions by promoting Pol II accumulation at the scnRNA production sites. It is worth noting that a similar phenomenon has been observed in worms: UAD-2 (an H3K27me3 reader containing IDRs) mediates liquid condensate formation, which is crucial for recruiting the piRNA transcriptional machinery to specific piRNA clusters and promoting piRNA precursor transcription. Disrupting condensate formation—either by IDR truncation or impairing SUMOylation—led to mislocalization of the piRNA transcriptional machinery to non-specific genomic regions (Zhu et al., 2025).

How could Rib1 mediate the accumulation of Pol II at the meiotic scnRNA production sites? Recent advances in eukaryotic transcriptional regulation have shown that the multivalent interactions of IDRs in Pol II, the Mediator complex, transcription factors, and some histone methyltransferases facilitate their aggregation at transcriptionally active loci (Cho et al., 2018; Dong et al., 2025; Namitz et al., 2023; Richter et al., 2022; Sabari et al., 2018). This aggregation enhances the local concentration of transcriptional machinery, thereby facilitating active transcription.

Several lines of evidence provided in this study suggest that Rib1 may promote Pol II aggregation in a similar way. First, three putative phase separation peptides were found within the intrinsically disordered N-terminal region of Rib1 (Figure 4B), and Rib1 forms aggregates in the MIC, especially at the telomere clustered regions (Figure 3A, (Tian et al., 2019)). Disruption of its aggregation, either by 1,6-hexanediol treatment or phase-separation peptides truncation, leads to an abnormally homogeneous distribution of Pol II on the MIC chromatin (Figure 4E-F). Second, restoration of Rib1 aggregation by fusion phase-separation peptides deleted Rib1 with a characterized phase-separation driving peptide also resulted in an accumulation of Pol II within the Rib1 condensates in the MIC (Figure 6F, Rib1-PSPΔ-FUS cells). The observation of unfunctional Rib1-PSPΔ-FUS in promoting the MIC ncRNA transcription may be due to either the Rib1 PSPs having additional roles that cannot be substituted by FUS, or aggregates formed by FUS were not favored for the MIC ncRNA transcription. Third, chromatin profiling data show that IDR deleted version of Rib1 (i.e., Rib1-CTD-NLS) exhibits a markedly reduced occupancy at scnRNA production sites, particularly in subtelomeric regions (Figure 5A). Correspondingly, such truncation led to a significantly crippled function of Rib1 in maintaining Pol II’s non-homogeneous localization in the MIC (Figure 4C).

Summing up, this study revealed that potential G-quadruplex-forming sequences with G/C-rich tracts are highly enriched in recent integrated TEs, they may act as signature motifs for initiating transcription of ncRNAs in the meiotic MIC (Figure 6G). Moreover, this study also elucidated that Pol II binds these sites in a Rib1-independent manner, yet its accumulation at these sites is facilitated by Rib1. Given that Rib1 is a Mediator complex-associated protein, we postulate that Rib1 may facilitate the accumulation of Mediator complex at the scnRNA production sites, and consequently, lead to the accumulation of Pol II at these sites (Figure 6G).

## Material and methods

### Strains and culture conditions

The inbred wild-type strains of *Tetrahymena thermophila* of different mating types, B2086 and CU428, were obtained from the American National *Tetrahymena* Stock Center at Cornell University (Table S1). The *rib1*Δ strains of different mating types were generated in a previous study (Tian et al., 2019). *Tetrahymena* cells were cultured in the modified Neff medium (0.25% [w/v] proteose peptone, 0.25% [w/v] yeast extract, 0.5% [w/v] glucose, and 33.3 mM FeCl_3_, all from Sigma-Aldrich, St. Louis, MO) at 30°C in an incubator, without shaking (Cassidy-Hanley et al., 1997). To induce conjugation, cells of different mating types were starved overnight in 10 mM Tris-HCl (pH 7.5), then equal amounts of starved cells were mixed.

Hexanediol treatment was performed as described in a previous study (Gao et al., 2024), with a minor modification. A final concentration of 4% (w/v) 1,6-hexanediol (Sangon Biotech, Shanghai) or 4% (w/v) 2,5-hexanediol (Aladdin Scientific, Shanghai) was added to the cells at the meiotic prophase I, when most of the MIC began to elongate.

### Small RNA sequencing and downstream bioinformatic analyses

The published WT meiotic small RNA sequencing data were utilized to identify the production sites of meiotic scnRNAs. Briefly, raw data were retrieved from the Sequence Read Archive under accession number SRR6814805 (Noto et al., 2015). Sequencing adapters were removed using Cutadapt (version 2.4,(Martin, 2011)). Small RNA reads shorter than 26 nt or longer than 32 nt were filtered out using Trim Galore (version 0.6.7). The processed data were then aligned to the MIC genome (https://www.ncbi.nlm.nih.gov/datasets/genome/GCA_016584475.1/) using Bowtie (version 1.3.1, (Langmead et al., 2009)). Uniquely mapped reads that did not align to rDNA, tRNA, snoRNA, or mtDNA were retained for downstream analysis. Signal tracks of small RNA sequencing read coverage were generated by processing the aligned small RNA-seq reads using deepTools2 (version 3.5.1, (Ramirez et al., 2016)). The sequencing reads were normalized using ‘bins per million mapped reads’ method. Regions with enriched scnRNA reads (i.e., scnRNA peaks) were identified using MACS3 (version 3.0.0, bdgpeakcall function, (Zhang et al., 2008)). Small RNA read coverage was visualized using either Integrative Genomics Viewer (version 2.18.4 (Robinson et al., 2011)) or pyGenomeTracks (version 3.8, (Lopez-Delisle et al., 2021)).

To examine small RNA by next-generation sequencing, total RNA from *rib1*Δ cells expressing Rib1-WT-HA or Rib1-NQΔ-HA was extracted from meiotic cells. The RNA was then used to generate small RNA sequencing libraries. Library preparation was conducted by Novogene (Novogene Technology, Tianjin), using the NEBNext^®^ Multiplex Small RNA Library Prep Set for Illumina® kit (New England Biolabs, Ipswich, MA), following the manufacturer’s instructions. Subsequently, library DNA with an insert size between 18 and 40 bp was selected, purified, and sequenced using the NovaSeq 6000 (Illumina, San Diego, CA). The resulting small RNA-seq data were analyzed similarly to the approach described above, with the only difference being the omission of the initial Cutadapt processing step. All small RNA-seq raw data generated in this study can be retrieved from the Sequence Read Archive (PRJNA1224863; https://dataview.ncbi.nlm.nih.gov/object/PRJNA1224863?reviewer=j6nqk3kqc7d2nf5ejkdprepvve).

Consensus DNA motifs shared by the meiotic scnRNA peaks were identified using RSAT (Santana-Garcia et al., 2022). Potential G-quadruplex-forming sequences (PQSs) were identified using G4Hunter with default settings (Brazda et al., 2019). The *Tetrahymena* Type A1 and Type B2 IES sequences were retrieved from the MIC supercontigs (version 2016, (Hamilton et al., 2016)), based on the published coordinates (Noto et al., 2015). Genome regions containing specific motif features (e.g., 5G and 5C) were identified using custom Python scripts.

### Protein sequence analysis and gene knock-in

The IUPred3 and PLAAC algorithms were used to identify intrinsically disordered regions and prion-like regions, respectively (Erdos et al., 2021; Lancaster et al., 2014). Putative phase-separation-driving peptides (PSPs) were identified using PSPhunter (Sun et al., 2024). Structural information for the Rib1 protein was predicted using AlphaFold and retrieved from UniProt (UniProt, 2025; Varadi et al., 2024). The protein 3D structure was visualized using PyMOL (version 2.6.0a0).

Gene knock-in was performed by introducing plasmid DNA into *Tetrahymena* cells. The plasmid DNAs used in this study are listed in Table S1. Below is a description of the generation of these plasmids:

To construct the plasmid DNA for generating *Tetrahymena* cells expressing the full-length Rib1 coding sequence from its endogenous locus, we first amplified the 5’ flanking sequence of *RIB1* and the entire *RIB1* coding sequence using primers #1 & #2. Primers were designed based on the latest gene model (Ye et al., 2025). Simultaneously, an 856 bp 3’ flanking sequence of *RIB1* was amplified using primers #3 & #4. Adapter sequences for ligation were added to the termini of the PCR products during amplification. Next, a DNA fragment containing the *EGFP* coding sequence, the *BTU1* terminator sequence, and the entire neo4 cassette was released from the pEGFP-neo4 plasmid (Kataoka et al., 2010) by BamHI-XhoI double digestion. The pBluescript II SK(+) vector backbone was also released from pEGFP-neo4 by SacI-KpnI double digestion. These four DNA fragments were then ligated by Gibson assembly (Gibson et al., 2009), using the NEBuilder® HiFi DNA Assembly Kit (New England Biolabs). The resulting plasmid is pRib1-WT-EGFP. To generate pRib1-WT-HA, the EGFP-BTU1 fragment was excised from pRib1-WT-EGFP using BamHI-PstI double digestion. A DNA fragment containing the Hemagglutinin tag (HA-tag) coding sequence and the *BTU1* terminator sequence was released from the pHA-neo4 plasmid (Kataoka et al., 2010) by BamHI-PstI digestion. The HA-BTU1 fragment was then ligated into the BamHI-PstI cut pRib1-WT-EGFP plasmid using T4 DNA ligase (Takara Bio, Kyoto), resulting in the plasmid pRib1-WT-HA. Plasmids used to generate strains expressing Rib1-NTD-HA, Rib1-CTD-HA, and Rib1-NQΔ-HA were constructed in a similar manner, using primers #5 - #10.

To generate the plasmid DNA used to express Rib1-NTD with a known MIC localization signal (Iwamoto et al., 2018), we started with pRib1-NTD-HA. First, the HA-tag coding sequence of pRib1-NTD-HA was excised using BamHI and SpeI. A synthetic DNA fragment (synthesized by Thermo Fisher Scientific, Waltham, MA; Frag1, Table S1) containing the MicNLS2 coding sequence, HA-tag coding sequence, and ligation adapter sequences was then integrated into the BamHI-SpeI cut pRib1-NTD-HA plasmid via Gibson assembly. The resulting plasmid is pRib1-NTD-NLS-HA. To generate pRib1-CTD-NLS-HA, the HA-BTU1 fragment was excised from pRib1-CTD-HA using BamHI and PstI. Meanwhile, the MicNLS2-HA-BTU1 DNA fragment was released from pRib1-NTD-NLS-HA with BamHI and PstI and ligated into the BamHI-PstI cut pRib1-CTD-HA using T4 DNA ligase, resulting in pRib1-CTD-NLS-HA.

The plasmid DNA used for expressing Rib1 with deletions of all PSPs was generated as follows: First, a backbone DNA was obtained by digesting the plasmid pBNMB1-CHA (Sequence can be retrieved from Zenodo; Link for reviewer: https://zenodo.org/records/14887167?preview=1&token=eyJhbGciOiJIUzUxMiJ9.eyJpZCI6IjA4YTQ4ZmE5LWE4YzAtNGVmMi04ZWNjLWM3NDA3YjQ1MTBjNyIsImRhdGEiOnt9LCJyYW5kb20iOiI3MjM5ZjIyOTc5ZmQ0Y2M5NDVhYWEyNmQ1NmQwMzcwMCJ9.v1KjBvr31iK3X2VWipNgFUIpRFtINbI-1OsQIL9LIqtNwv2YCBFlCC7Qf25iNmGF1ADNDU3GPd9ZW_DxHTzO7g) with NdeI and BamHI. A DNA fragment containing ligation adapters and the N-terminal part of the PSP-deleted *RIB1* coding sequence was synthesized by Tsingke (Tsingke Biology, Beijing; Frag2, Table S1). The C-terminal *RIB1* coding sequence was then amplified from *Tetrahymena thermophila* genomic DNA using primers #11 and #12, with adapter sequences fused to the termini. The three DNA fragments were ligated by Gibson assembly, resulting in pRib1-PSPΔ-HA.

Plasmids expressing specific PSP-deleted Rib1 variants were constructed similarly, with variations in the *RIB1* coding sequence preparation. For example, the Rib1-PSPaΔ-HA coding sequence was assembled using two DNA fragments amplified from the genomic DNA with primers #13 & #14 and #12 & #15. Similarly, Rib1-PSPbΔ-HA and Rib1-PSPcΔ-HA coding sequences were constructed using different combinations of primers (#13 & #16, #12 & #17, #13 & #18, and #12 & #19). To add the MicNLS2 sequence to these plasmids, the HA-tag was replaced with a MicNLS2-HA-tag coding sequence. As an example, for pRib1-PSPΔ-NLS-HA, the HA-tag of pRib1-PSPΔ-HA was removed by BamHI-SpeI double digestion, and the MicNLS2-HA-tag coding sequence was amplified from the pRib1-CTD-NLS-HA plasmid using primers #20 & #21. These fragments were then ligated by Gibson assembly, resulting in the plasmid pRib1-PSPΔ-NLS-HA.

To generate the plasmid DNA used for expressing truncated Rib1 fused with the phase separation-driving region of FUS protein (hereinafter referred to as FUS; (Murthy et al., 2019)), we excised the Rib1 coding sequence from the pRib1-PSPΔ-HA plasmid using NdeI-BamHI double digestion. A DNA fragment containing part of the Rib1 N-terminal coding sequence and the *FUS* coding sequence was synthesized by Sangon Biotech (Frag3, Table S1). The remaining PSP-deleted *RIB1* coding sequence was amplified from pRib1-PSPΔ-HA using primers #12 & #15. These three DNA fragments were ligated by Gibson assembly, resulting in the plasmid pRib1-PSPΔ-FUS-HA.

To determine whether the FUS peptide alone could drive Pol II accumulation, we generated an EGFP-tagged FUS construct. The backbone vector was prepared by digesting pBNMB1-EGFP with NdeI and BamHI. Meanwhile, the FUS coding sequence along with its flanking adapters was amplified from pRib1-PSP Δ -FUS-HA, using primers #22 & #23. Then, these two DNA fragments were ligated by Gibson assembly, resulting in the plasmid pEGFP-FUS.

The biolistic transformation method was used to introduce the above plasmid DNA into starved *rib1*Δ cells, using GJ-1000 (SCIENTZ, Ningbo, China) (Hao et al., 2024), following a published protocol (Cassidy-Hanley et al., 1997). To facilitate the integration of recombinant DNA into the *Tetrahymena* MAC genome, the plasmid DNA used for transformation was linearized by SacI or KpnI digestion prior to transformation. After transformation, recombinant DNA-containing transformants were selected by culturing in modified Neff medium supplemented with 120 μg/ml paromomycin (Mochizuki, 2008).

### Immunostaining, in situ hybridization, and immunoblotting analyses

Protein localization was determined by indirect immunostaining as previously described (Loidl et al., 2012; Tian et al., 2022), with minor modifications. Briefly, approximately 1 × 10⁶ cells were fixed and permeabilized at 25°C for 30 minutes in a solution containing 3.7% (v/v) formaldehyde and 0.5% (v/v) Triton X-100. The fixed cells were then harvested and resuspended in 500 µl of ice-cold fixation solution containing 4% (v/v) formaldehyde and 3.4% (w/v) sucrose. The cells were subsequently spread onto a slide and air-dried for 2 hours at 25°C. To immunostain the protein of interest, air-dried slides were first rehydrated in IF buffer (1 × PBS containing 0.1% [v/v] Tween-20), followed by incubation with blocking buffer (0.1% [v/v] Tween-20, 3% [w/v] Bovine Serum Albumin, 10% [v/v] normal goat serum, and 1 × PBS). Primary antibodies, resuspended in the blocking buffer, were applied to the slides, which were then incubated at 25°C for 2 hours or at 4°C overnight. The HA-tagged protein was detected using either anti-HA rabbit monoclonal antibody (Clone C29F4, 1:500 dilution, Cell Signaling Technology, Danvers, MA) or anti-HA mouse monoclonal antibody (Clone 6E2, 1:200 dilution, Cell Signaling Technology). *Tetrahymena* Rpb3 was detected using a custom antibody (Kataoka and Mochizuki, 2017). After primary antibody incubation, the slides were rinsed with the IF buffer. Then, appropriate fluorescence-conjugated secondary antibodies that resuspended in the blocking buffer (1:1000 dilution) were used to label the primary antibodies. Double-stranded RNA staining was performed as previously described, using the J2 antibody (Nordic-MUbio, Fargo, ND; (Tian et al., 2019; Woo et al., 2016)).

DNA elimination was investigated by labeling *REP2* IESs in *Tetrahymena* cells harvested 32 hours after induction of conjugation. Fluorescence *in situ* hybridization (FISH) labeling of *REP2* IESs was conducted as previously described, without modification (Noto et al., 2010). The DNA fragment used for preparing the FISH probe was amplified from *Tetrahymena* genomic DNA using primers #24 & #25 (Table S1). The stacked images captured using a Zeiss Z2 microscope (Carl Zeiss, Thornwood, NY) were subsequently processed through deconvolution, merging, and conversion into 2D images using FIJI software (Schindelin et al., 2012). Since a monochrome camera was used for imaging, the grayscale fluorescence signals captured from different channels were colored using Photoshop (version CS5, Adobe, San Jose, CA). For each slide, around 50 cells were examined. In cases where all cells showed consistent patterns, a representative image was selected and included in the figure.

To detect *Tetrahymena* proteins by Western Blotting, 1.5 × 10⁶ cells of interest were washed once with 10 mM Tris-HCl (pH 7.4), and crude proteins were precipitated using 10% (v/v) Trichloroacetic acid. After incubating the cells with Trichloroacetic acid on ice for 10 minutes, the crude protein pellet was collected by centrifugation (5,000 rpm, 1 minute at 4°C, using Eppendorf Centrifuge 5430R). The pellet was resuspended in 1x LDS buffer (Thermo Fisher Scientific, Waltham, MA), denatured at 70°C, and then loaded onto an SDS-PAGE gel for electrophoresis. Separated proteins were transferred onto a PVDF membrane (GE Amersham, Fairfield County, CT) using the eBlot^®^ L1 system (GenScript Biotech Corporation, Piscataway, NJ), following the manufacturer’s instructions. Proteins on the membrane were detected using anti-HA rabbit monoclonal antibody (Clone C29F4, 1:2,000 dilution, Cell Signaling Technology), HRP-conjugated secondary antibodies, and Clarity™ Western ECL Substrate (Bio-Rad Laboratories, Hercules, CA). Visualization and documentation of immunoblot membranes were carried out using the ChemiDoc^TM^ Imaging System (Bio-Rad Laboratories).

### Chromatin profiling using CUT&Tag

To elucidate the binding sites of Rpb3 on the meiotic MIC chromatin, approximately 3 × 10^8^ cells expressing Rpb3-HA from its endogenous promoter were starved and mixed to induce meiosis. After confirming that over 90% of the cells were in meiotic prophase I (approximately 2.5 hours post-mixing), with the MICs showing slight elongation, cells were harvested, lysed, and MICs were isolated following a previously established protocol (Gorovsky, 1970). After purification, approximately 5 × 10^6^ MICs, with a purity of over 99.9%, were used for CUT&Tag analysis.

Key commercial reagents used in the CUT&Tag experiment are listed in Table S1. To enhance the binding of purified MICs to Concanavalin A beads, MICs were first equilibrated in a buffer containing 150 mM NaCl, 0.5 mM Spermidine, 0.1% (w/v) Bovine Serum Albumin, and 10 mM HEPES (4-[2-hydroxyethyl]-1-piperazineethanesulfonic acid, pH 7.4) for 5 minutes. CUT&Tag was then performed using BioMag^®^Plus Concanavalin A beads (Bangs Laboratories, Fishers, IN), the Hyperactive^®^ Universal CUT&Tag Assay Kit for Illumina (Vazyme, Nanjing), the anti-HA rabbit monoclonal antibody (Clone C29F4), and the complementary secondary antibody. As a control, an equal number of MICs isolated from untagged WT meiotic cells were processed in parallel. CUT&Tag libraries were constructed using indexed primers (Buenrostro et al., 2015) and the KAPA HiFi HotStart Real-Time Library Amplification Kit (Roche). The libraries were subsequently sequenced on NovaSeq 6000 (Illumina).

After obtaining the raw sequencing data, low-quality reads and sequencing adapters were removed using Trim Galore (version 0.6.7). The remaining reads were aligned to the MIC genome (https://www.ncbi.nlm.nih.gov/datasets/genome/GCA_016584475.1/) using Bowtie 2 (version 2.3.5.1, (Langmead and Salzberg, 2012)). PCR duplicates were removed using Picard (2.25.5). Reads that aligned to multiple genomic sites, rRNAs, tRNAs, snoRNAs, or mitochondrial DNA sequences were discarded. The remaining uniquely mapped reads were used for downstream analyses. Normalized CUT&Tag read coverage profiles were generated using deepTools2 (3.5.1). The embedded normalization method (reads per genome content) was employed for normalization of CUT&Tag data. Aberrant Rpb3 binding sites in *rib1*Δ MICs were identified by comparing Rpb3 CUT&Tag data from *rib1*Δ and WT MICs using MACS3 (3.0.0). Consensus DNA motifs overrepresented at these aberrant sites were identified using STREME (Bailey, 2021). Coverage plots of sequencing reads around the regions/sites of interest were generated using deepTools2 (3.5.1). All CUT&Tag raw data generated in this study can be retrieved from the Sequence Read Archive (PRJNA1224863; https://dataview.ncbi.nlm.nih.gov/object/PRJNA1224863?reviewer=j6nqk3kqc7d2nf5ejkdprepvve).

## Compliance and ethics

The author(s) declare that they have no conflict of interest.

## Acknowledgements

This work is supported by the Science & Technology Innovation Project of Laoshan Laboratory (LSKJ202203202), the National Natural Science Foundation of China (32370451), the Fundamental Research Funds for the Central Universities (202241003), the Natural Science Foundation of Shandong Province (ZR2022JQ13), and by the European Union’s Horizon 2020 research and innovation programme under the Marie Skłodowska-Curie grant agreement (101024333) to M.T., Labex EpiGenMed Advanced Grant from Agence Nationale de la Recherche (ANR-10-LABX-12-01) and Equipes 2022 Grant from Fondation pour la Recherche Médicale (EQU202203014651) to K.M. We thank Dr. Yongqiang Liu, Dr Feng Gao, and Dr. Yunyi Gao (Ocean University of China) for their technical supports. We thank Dr. Kai Chen (Southern Marine Science and Engineering Guangdong Laboratory [Guangzhou], China) for sharing critical reagents with us. We appreciate the computing resources provided by IEMB-1, a high-performance computing cluster operated by the Institute of Evolution & Marine Biodiversity. We also acknowledge the support of the High-Performance Biological Supercomputing Center at the Ocean University of China for this research.

