## Supplemental figure 1-4 for "Transposon targeting non-coding RNA transcription targets G/C-rich tracts and is facilitated by an intrinsically disordered protein in *Tetrahymena*"

### Figure S1

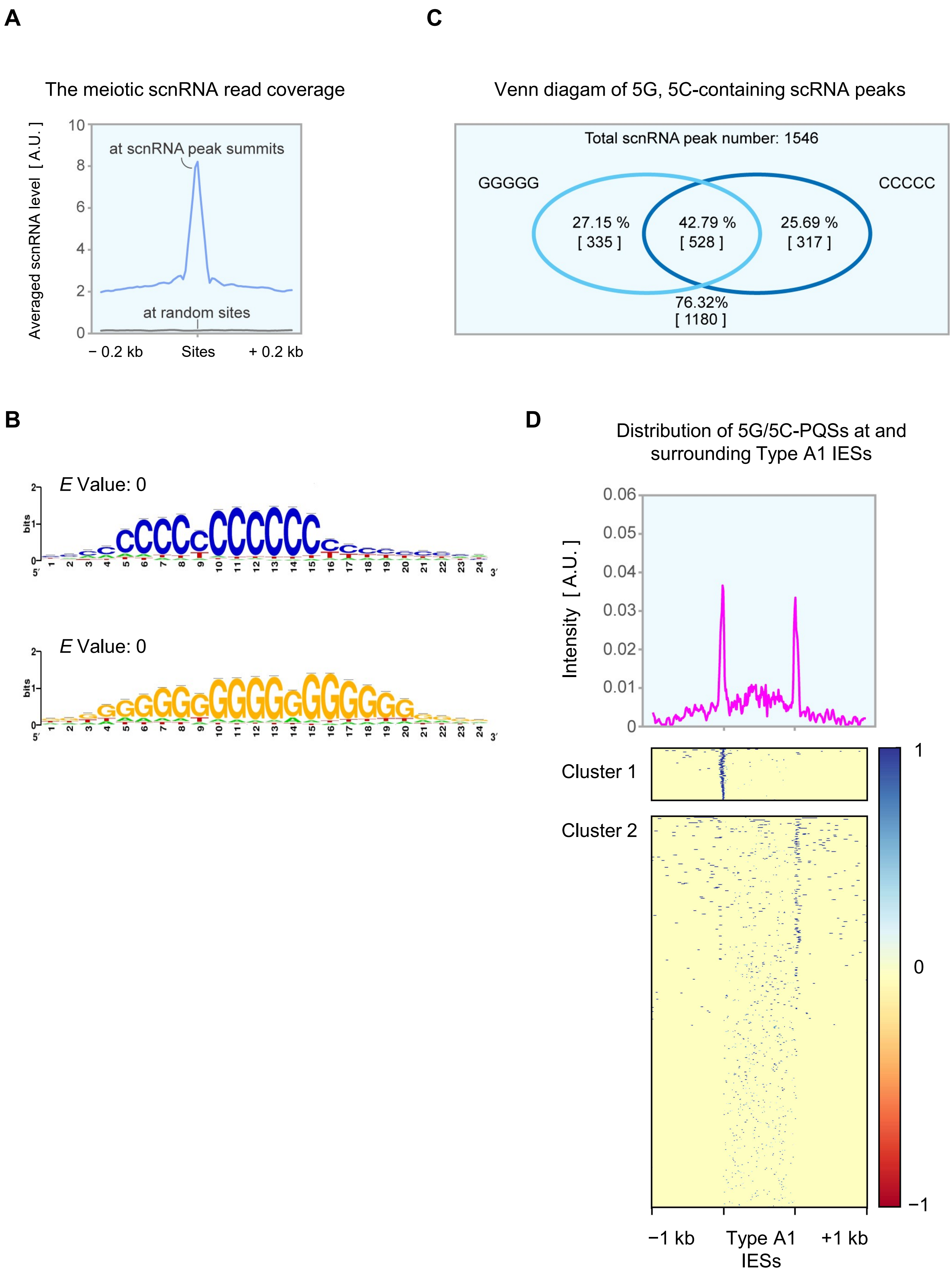
A
The meiotic scnRNA read coverage
Averaged scnRNA level [ A.U. ]
10
8
6
4
2
0
at scnRNA peak summits
at random sites
− 0.2 kb
Sites
+ 0.2 kb
B
E Value: 0
bits
E Value: 0
bits
5′
3′
C
Venn diagam of 5G, 5C-containing scRNA peaks
Total scnRNA peak number: 1546
GGGGG
CCCCC
27.15 %
[ 335 ]
42.79 %
[ 528 ]
25.69 %
[ 317 ]
76.32%
[ 1180 ]
D
Distribution of 5G/5C-PQSs at and surrounding Type A1 IESs
Intensity [ A.U. ]
0.06
0.05
0.04
0.03
0.02
0.01
0
Cluster 1
Cluster 2
1
0
−1
−1 kb
Type A1 IESs
+1 kb

**Figure S1. Enrichment of G/C-rich tracts containing G-quadruplex-forming sequences in the MIC ncRNA transcription regions. Related to Figure 1.**

A) Visualization of averaged scnRNA read coverage at and surrounding scnRNA peak summits compared to random sites. The colorimetric scale (red to blue) represents coverage intensity, with red indicating higher enrichment.
B) Venn diagram illustrating the overlap and statistical analysis of scnRNA peaks containing GGGGG (5G) and CCCCC (5C) tracts.
C) Significantly enriched consensus DNA motifs identified from scnRNA peak sequences using RSAT.
D) Upper panel: Distribution of 5G/5C-containing putative G-quadruplex-forming sequences (5G/5C-PQSs) at and around Type A1 IESs. Lower panels: Heatmap visualize relative positions of 5G/5C-PQSs at each investigated regions. Short blue lines denote regions with 5G/5C-PQSs. Clustering analysis was performed using the k-means method provided by DeepTools2. The colorimetric scale (blue to red) represents coverage intensity, with blue indicating higher enrichment. These results indicate that 5G/5C-PQSs are preferentially enriched at a single terminus of some Type A1 IESs, but not at both termini. The biological significance of this asymmetric distribution remains unclear.

### Figure S2

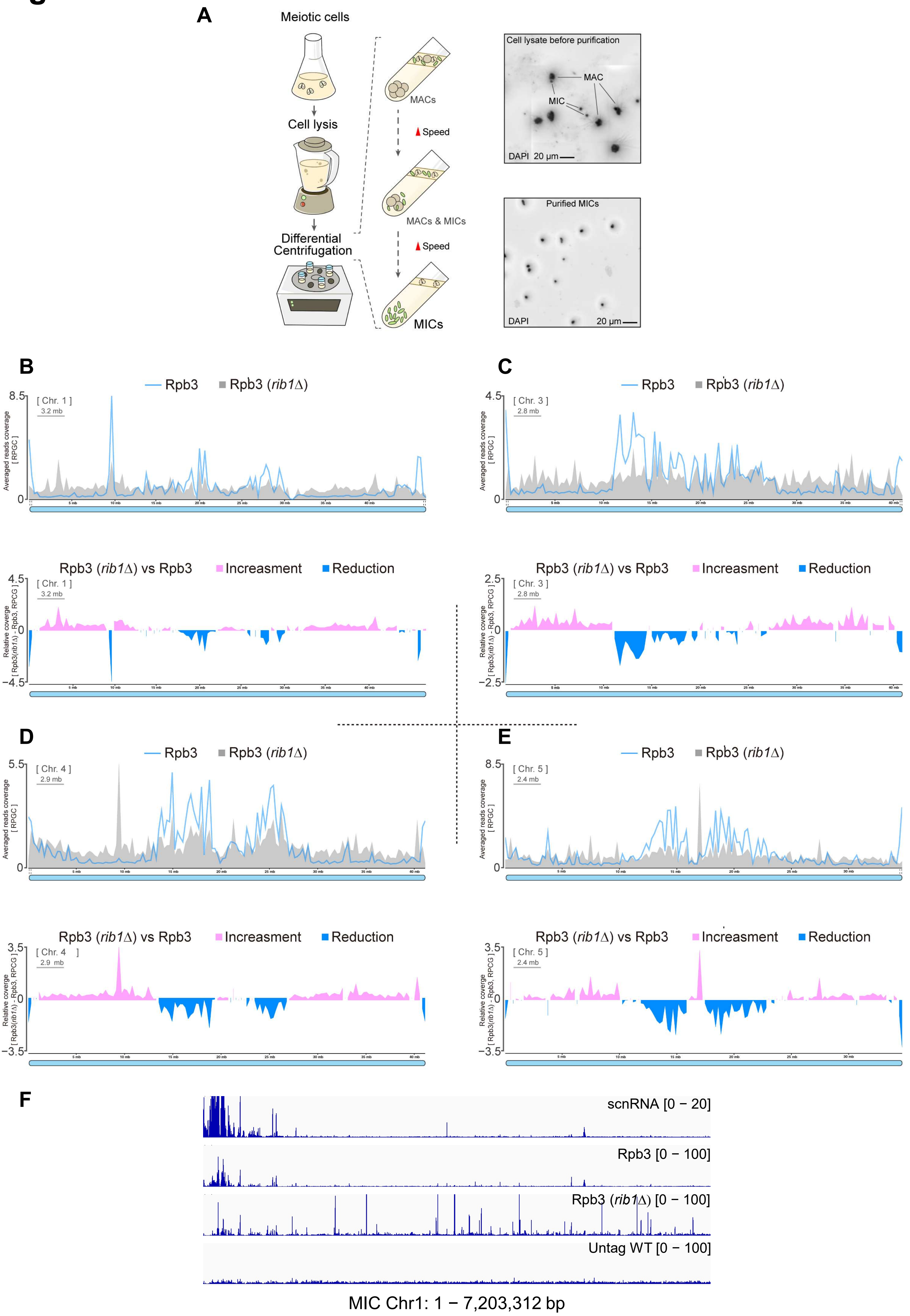

A
Meiotic cells
Cell lysis
Differential Centrifugation
MACs
Speed
MACs & MICs
Speed
MICs
Cell lysate before purification
MAC
MIC
DAPI 20 µm
Purified MICs
DAPI
20 µm
B
Rpb3
Rpb3 (rib1Δ)
[ Chr. 1 ]
3.2 mb
Averaged reads coverage [ RPGC ]
Rpb3 (rib1Δ) vs Rpb3
Increasment
Reduction
Relative coverge [ Rpb3(rib1Δ) - Rpb3, RPCG ]
C
[ Chr. 3 ]
2.8 mb
D
[ Chr. 4 ]
2.9 mb
E
[ Chr. 5 ]
2.4 mb
F
scnRNA [0 − 20]
Rpb3 [0 − 100]
Rpb3 (rib1Δ) [0 − 100]
Untag WT [0 − 100]
MIC Chr1: 1 − 7,203,312 bp

**Figure S2. Chromatin profiling analysis of RNA polymerase II-specific subunit in the MIC using CUT&Tag. Related to Figure 2.**

A) Left panels: Step-by-step workflow of the MIC isolation procedure, highlighting key stages such as cell lysis, purification, and enrichment of MICs; Right panels: Representative microscopy images showing the isolated MICs. Scale bars: 20 μm.
B-E) Upper panels: Normalized Rpb3-HA CUT&Tag read coverage in WT or *rib1*Δ MIC chromosomes. Lower panels: Relative CUT&Tag read coverage showing the increase and decrease of Rpb3 occupancy on MIC chromosomes in response to *RIB1* deletion.
F) The scnRNA sequencing read coverage and the normalized Rpb3-HA CUT&Tag read coverage on the left end of MIC chromosome 1 in WT and *rib1*Δ cells are shown. CUT&Tag read coverage obtained from untagged WT MICs is included as a negative control.

### Figure S3

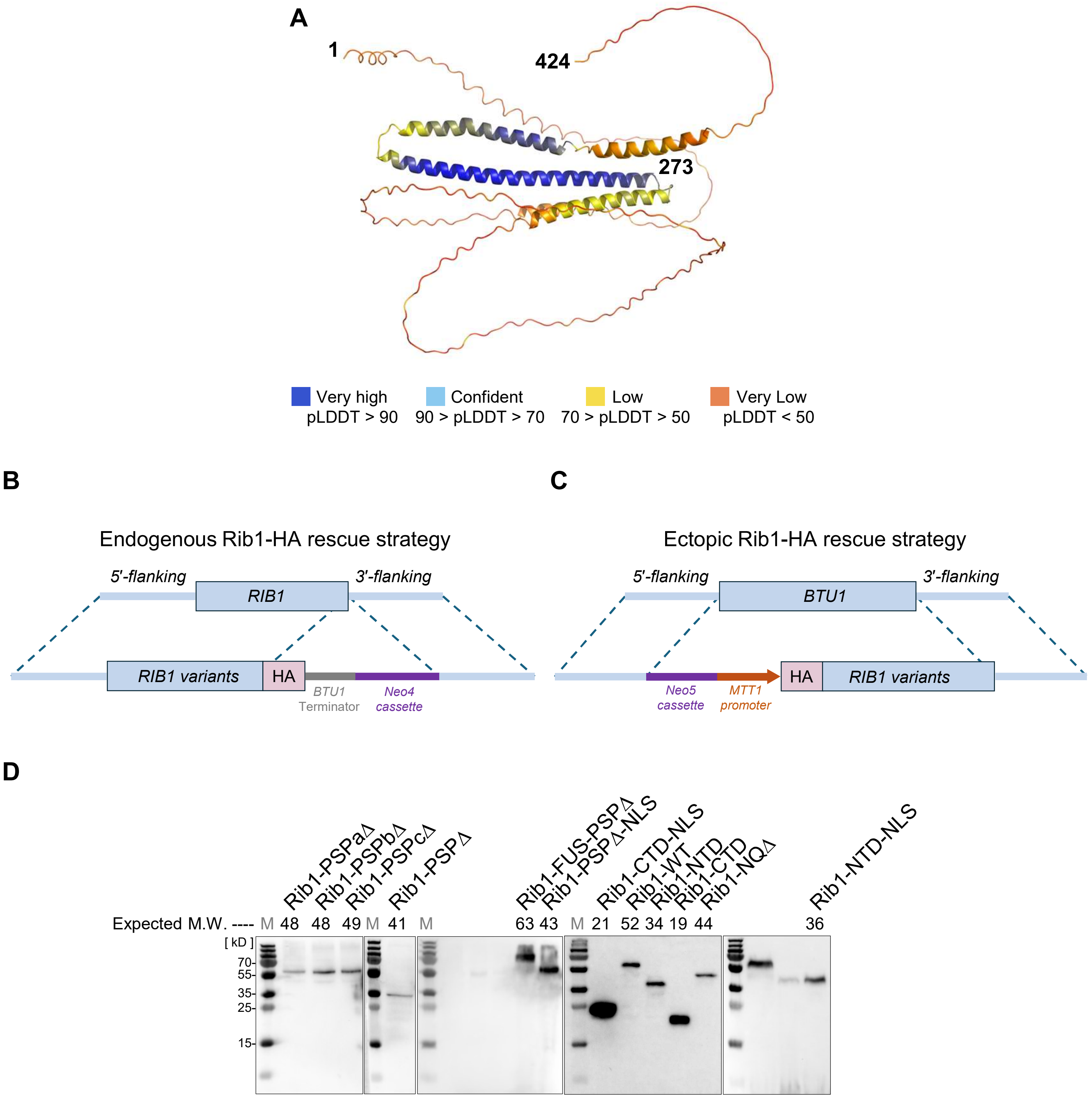


**Figure S3. Predicted Rib1 structure and Rib1-tagging strategies. Related to Figure 4.**

A) Predicted three-dimensional structure of Rib1 generated by AlphaFold. The protein is color-coded based on the per-residue prediction confidence score (pLDDT), with blue indicating high confidence and red indicating low confidence.
B-C) Schematic diagrams of the Rib1 ectopic tagging strategies, highlighting the insertion sites for epitope tags.
D) Confirmation of protein expression using western blotting.

### Figure S4

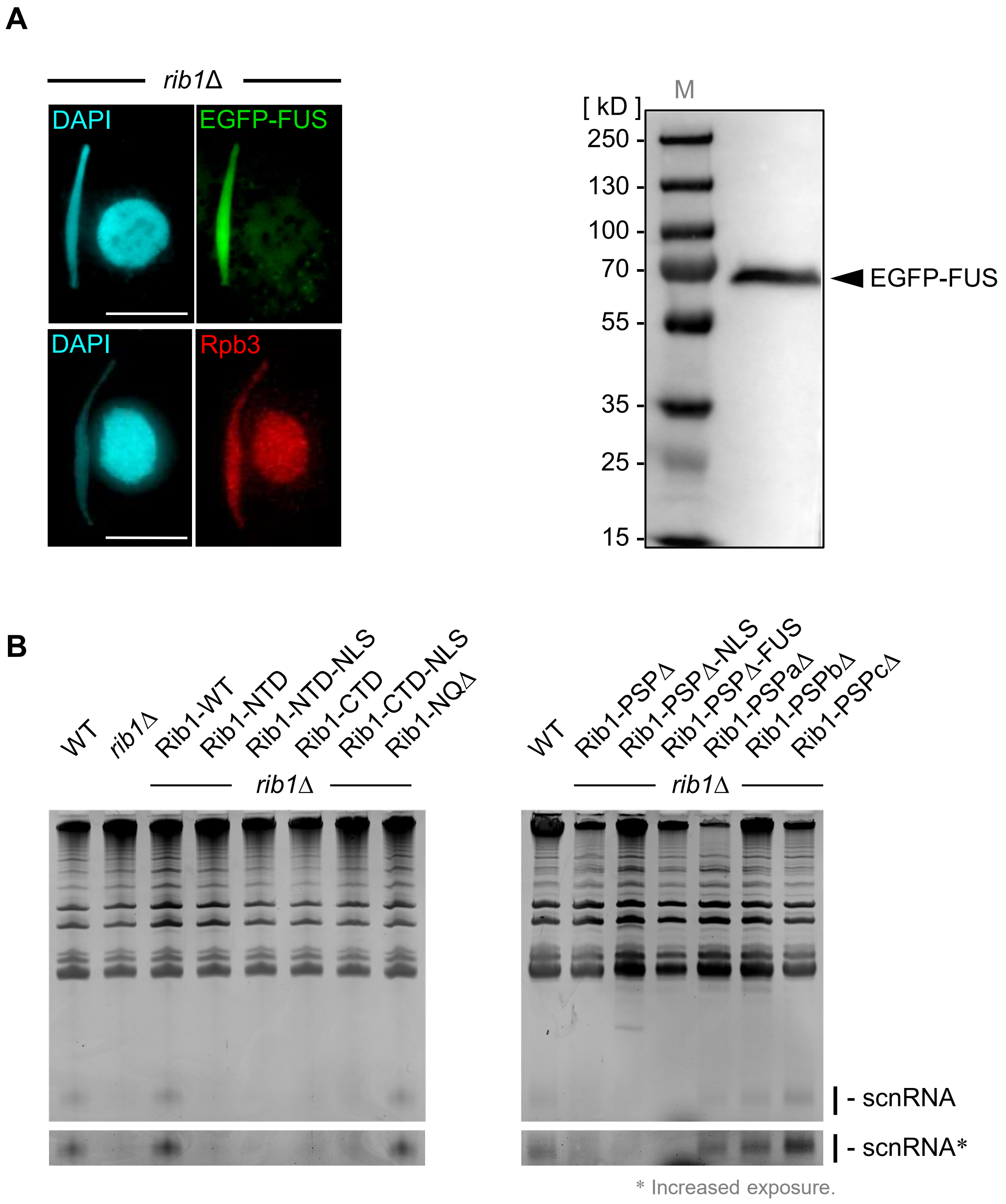


**Figure S4. Supplementary images related to Figure 4.**

A) Left panels: Localizations of EGFP-FUS and Rpb3 in *rib1*Δ cells expressing EGFP-FUS. Scale bars: 10 µm. Right panel: Immunoblotting of analysis of *rib1*Δ cells expressing EGFP-FUS.

B) Uncropped scnRNA gel electrophoresis images, related to Figure 4D.
