## Supplemental Table 1 for "Transposon targeting non-coding RNA transcription targets G/C-rich tracts and is facilitated by an intrinsically disordered protein in *Tetrahymena*"

**Table S1. Key resources table.**

| Categories | IDs | Oligo or DNA fragment IDs | Source | Sequence (5'->3') |
|---|---|---|---|---|
| *Tetrahymena* strains | *T. thermophila:* B2086 II, mat1-2/mat1-2 (mat1-2; II) | N/A | *Tetrahymena* Stock Center | N/A |
| | *T. thermophila:* CU428.2, mpr1-1/mpr1-1 (MPR1; mp-s, VII) | N/A | *Tetrahymena* Stock Center | N/A |
| | *T. thermophila: rib1* Δ B2086 | N/A | Tian et al., 2019 | N/A |
| | *T. thermophila: rib1* Δ CU428 | N/A | Tian et al., 2019 | N/A |
| | *T. thermophila* : *rib1* Δ B2086/Rib1-CTD-NLS-HA | N/A | This study | N/A |
| | *T. thermophila* : *rib1* Δ CU428/Rib1-CTD-NLS-HA | N/A | This study | N/A |
| | *T. thermophila* : *rib1* Δ B2086/Rib1-WT-HA | N/A | This study | N/A |
| | *T. thermophila* : *rib1* Δ CU428/Rib1-WT-HA | N/A | This study | N/A |
| | *T. thermophila* : *rib1* Δ B2086/Rib1-NTD-HA | N/A | This study | N/A |
| | *T. thermophila* : *rib1* Δ CU428/Rib1-NTD-HA | N/A | This study | N/A |
| | *T. thermophila* : *rib1* Δ B2086/Rib1-NTD-NLS-HA | N/A | This study | N/A |
| | *T. thermophila* : *rib1* Δ CU428/Rib1-NTD-NLS-HA | N/A | This study | N/A |
| | *T. thermophila* : *rib1* Δ B2086/Rib1-CTD-HA | N/A | This study | N/A |
| | *T. thermophila* : *rib1* Δ CU428/Rib1-CTD-HA | N/A | This study | N/A |
| | *T. thermophila* : *rib1* Δ B2086/Rib1-NQΔ-HA | N/A | This study | N/A |
| | *T. thermophila* : *rib1* Δ CU428/Rib1-NQΔ-HA | N/A | This study | N/A |
| | *T. thermophila* : *rib1* Δ B2086/Rib1-PSPΔ-HA | N/A | This study | N/A |
| | *T. thermophila* : *rib1* Δ CU428/Rib1-PSPΔ-HA | N/A | This study | N/A |
| | *T. thermophila* : *rib1* Δ B2086/Rib1-PSPaΔ-HA | N/A | This study | N/A |
| | *T. thermophila* : *rib1* Δ CU428/Rib1-PSPaΔ-HA | N/A | This study | N/A |
| | *T. thermophila* : *rib1* Δ B2086/Rib1-PSPbΔ-HA | N/A | This study | N/A |
| | *T. thermophila* : *rib1* Δ CU428/Rib1-PSPbΔ-HA | N/A | This study | N/A |
| | *T. thermophila* : *rib1* Δ B2086/Rib1-PSPcΔ-HA | N/A | This study | N/A |
| | *T. thermophila* : *rib1* Δ CU428/Rib1-PSPcΔ-HA | N/A | This study | N/A |
| | *T. thermophila* : *rib1* Δ B2086/Rib1-PSPΔ-NLS-HA | N/A | This study | N/A |
| | *T. thermophila* : *rib1* Δ CU428/Rib1-PSPΔ-NLS-HA | N/A | This study | N/A |
| | *T. thermophila* : *rib1* Δ B2086/Rib1-PSPΔ-FUS-HA | N/A | This study | N/A |
| | *T. thermophila* : *rib1* Δ CU428/Rib1-PSPΔ-FUS-HA | N/A | This study | N/A |
| | *T. thermophila* : *rib1* Δ B2086/EGFP-FUS | N/A | This study | N/A |
| | *T. thermophila* : *rib1* Δ CU428/EGFP-FUS | N/A | This study | N/A |
| | *T. thermophila* : Rpb3-HA B2086 | N/A | Tian et al., 2019 | N/A |
| | *T. thermophila* : Rpb3-HA CU428 | N/A | Tian et al., 2019 | N/A |
| | *T. thermophila* : *rib1* Δ B2086/Rpb3-HA | N/A | This study | N/A |
| | *T. thermophila* : *rib1* Δ CU428/Rpb3-HA | N/A | This study | N/A |
| | *T. thermophila* : Rib1-HA B2086 | N/A | Tian et al., 2019 | N/A |
| | *T. thermophila* : Rib1-HA CU428 | N/A | Tian et al., 2019 | N/A |
| Plasmids | pEGFP-neo4 | N/A | Kataoka et al., 2010 | N/A |
| | pHA-neo4 | N/A | Kataoka et al., 2010 | N/A |
| | pRib1-WT-EGFP-neo4 | N/A | This Study | N/A |
| | pRib1-WT-HA-neo4 | N/A | This Study | N/A |
| | pRib1-NTD-HA-neo4 | N/A | This Study | N/A |
| | pRib1-CTD-HA-neo4 | N/A | This Study | N/A |
| | pRib1-NQΔ-HA-neo4 | N/A | This Study | N/A |
| | pRib1-NTD-NLS-HA-neo4 | N/A | This Study | N/A |
| | pRib1-CTD-NLS-HA-neo4 | N/A | This Study | N/A |
| | pBNMB1-CHA-neo5 | N/A | https://zenodo.org/records/148 | N/A |
| | pRib1-PSPΔ-HA-neo5 | N/A | This Study | N/A |
| | pRib1-PSPaΔ-HA-neo5 | N/A | This Study | N/A |
| | pRib1-PSPbΔ-HA-neo5 | N/A | This Study | N/A |
| | pRib1-PSPcΔ-HA-neo5 | N/A | This Study | N/A |
| | pRib1-PSPΔ-NLS-HA-neo5 | N/A | This Study | N/A |
| | pRib1-PSPΔ-FUS-HA-neo5 | N/A | This Study | N/A |
| | pEGFP-FUS-neo5 | N/A | This Study | N/A |
| | 1 | Rib1_5f964_SacG | This Study | ACTAAAGGGAACAAAAGCTGGAGCTCtgtgttggcttcttttatgaa |
| | 2 | Rib1_CDSr3027_BamH1G | This Study | TCTTCACCCTTAGAAACCATGGATCCGCTTTGGATATTTTCACAACAG |
| | 3 | Rib1_3f3049_Xho1G | This Study | GCTTATCGATACCGTCGACCTCGAGACAATCTTAGTAAGGAGCATTTATC |
| | 4 | Rib1_3r3854_Kpn1Gib | This Study | CTCACTATAGGGCGAATTGGGTACCAATAGCTGTGAATCTTTTACCG |
| | 5 | Rib1_1-272_rv1 | This Study | TTCTTCACCCTTAGAAACCATGGATCCttaatcgaagtatctttctttatcag |
| | 6 | Rib1_5utrNQTRr2201 | This Study | CATTTAAAAAAAATCTTTTTAAAGT |
| | 7 | Rib1_273fv2 | This Study | tagagtgactttaaaaagattttttttaaatggatagaaagatcaagtaaattgtagaac |
| | 8 | Rib1_84-142d_Rv1 | This Study | agaattttaataaccagactcaggcatc |
| | 9 | Rib1_cdsNQtrF2182 | This Study | ATTCCTGGCTAGAATCCAAAT |
| | 10 | Rib1_84-142d_fv1 | This Study | gatgcctgagtctggttattaaaattctattcctggctagaatccaaat |
| | 11 | Rib1-CDS-down_f5924 | This Study | ATAATCAGTATATGCAGAATAAGTTC |
| | 12 | Rib1-CDS-down_r6536_SpeI | This Study | AGTAGCATAATCAGGAACATCATAAGGATAGGATCCGCTTTGGATATTTTCACAACAG |
| | 13 | Rib1_CDS_f5470_NdeI | This Study | TAATAAAATAAATAATAATACTAAACTTAAACATATGCAAGAAACAAATAATTTGTAG |
| | 14 | Rib1_CDS_PSP-A_r5675 | This Study | TATCTAGTTATAATTTGTATTAATCTAATTAGAAAGATTCTTATTGTTGTTTTACCTG |
| | 15 | Rib1_CDS_PSP-A_f5641 | This Study | TCTAATTAGATTAATACAAATTATAACTAGATAAATTAG |
| | 16 | Rib1_CDS_PSP-B_r5933 | This Study | TCTACAATAGTCCTTTTAGAATTTATAAGACATTTAATTTTAATTCTAGTTCATCGG |

| | | | | |
|---|---|---|---|---|
| Synthesized oligos and DNA fragments | 17 | Rib1_CDS_PSP-B_f5902 | This Study | TCTTATAAATTCTAAAAGGACTATTGTAGAC |
| | 18 | Rib1_CDS_PSP-C_r6153 | This Study | AGAACTTATTCTGCATATACTGATTATCGTAATAGAAAAATTCTCCTCTAGTAGGATTC |
| | 19 | Rib1_CDS_PSP-C_f6119 | This Study | TATTACGATAATCAGTATATGCAGAATAAG |
| | 20 | MicNLS2-HA_f6476_BamHI | This Study | ACTTTCTGTTGTGAAAATATCCAAAGCGGATCCAAGGGTAAGAAAAAATCCAAAGAAGG |
| | 21 | MicNLS2-HA_r6634_SpeI | This Study | ATTGGTTGGTTTAGCTGACCGATTCAGTTCGCTCAACTAGTAGCATAATCAGGAACATC |
| | 22 | hsFUS_f6116_EGFP | This Study | TTGGTATGGATGAATTATATAAGGGATCCGCTTCTAATGATTATACTTAATAAGCTAC |
| | 23 | hsFUS_r6819_EGFP | This Study | TTGGTTTAGCTGACCGATTCAGTTCGCTCAACTAGTACCTCTATCCTGTTATCCATAAC |
| | 24 | REP2FW | Noto et al., 2010 | TTGATGACTTAGATGACATTGATGAC |
| | 25 | REP2RV | Noto et al., 2010 | ACATTTCCAGCAGAATTGTCCAGC |
| | Frag1 | Frag1 | This Study | gataaagaaagatacttcgattaaGGATCCAAGGGTAAGAAAAAATCCAAAGAAGGAAAGACTGGAGCTTATGGCAAGAAGGCAAATTATCCTTATGATGTTCCTGATTATGCTACTAGTTGAGCGAACTGAATCGGTCAGCTAA |
| | Frag2 | Frag2 | This Study | CTAATAAAATAAATAATAATACTAAACTTAAACATCATATGATGCAAGAAACAAATAATTTGTAGTAGATCTTGAGTTCAAATAACACCAAAAAGCCTAATAATCCTCAGTCTTAACAGTAAATCTAGAGCACCTCTTAGTTTTTCAACAAGAACAGGTAAAACAACAATAAGAATCTTTCTAATTAGATTAATACAAATTATAACTAGATAAATTAGAATTAGATGAACCAAAATCCAAACTAGATCAATCAAAATCCTAATTAAATGAACTAAAACCAAAATCAGATGAACTAGAATCAAAATCCGATGAACTAGAATTAAAATTAAATGTCTTATAAATTCTAAAAGGACTATTGTAGACCTTAAATGAATATGCAAAACAACTTTTAAAGCTATCCCAATAACAGAAATTACGAAAATAATATGAATCCTACTAGAGGAGAATTTTTCTATTACGATAATCAGTATATGCAGAATAAGTTC |
| | Frag3 | Frag3 | This Study | CTAATAAAATAAATAATAATACTAAACTTAAACATCATATGATGCAAGAAACAAATAATTTGTAGTAGATCTTGAGTTCAAATAACACCAAAAAGCCTAATAATCCTCAGTCTTAACAGTAAATCTAGAGCACCTCTTAGTTTTTCAACAAGAACAGGTAAAACAACAATAAGAATCTTgcttctaatgattatacttaataagctacttaatcttatggtgcttatcctacttaacctgggcagggttattcgtagtaatccagccaaccttatggccagtaatcctattcgggttattcttaatctactgatacttcgggatatgggcagtccagctattcaagctatggctaatctcagaatactggttatggtacgtagagcaccccgcagggctatggctcgactggcgggtacggctcgtcctagagctctcagtcctcttacggtcagtagtcttcgtaccctggttatggttaataacctgctccttcttctacttctggttcgtacgggtcttcctcacagtcgagcagttacggttagcctcagtctggttcttactcccagtagcctagctacggcggtcagtaacaaagctacggtcagtagtaaagctacaatcctcctcagggttacggttagtaaaattaatacaatagctcttccggtggcggaggaggcggcggagggggtggaaattacggccaggatcaaagctccatgagcagtggtggcggttccggaggtggctatggtaatcaggatcagtctggtggcggtggatctggcggttatggataacaggatagaggtTCTAATTAGATTAATACAAATTATAACTAGATAAATTAG |
